# Modeling mitochondrial inheritance enables high-precision single-cell lineage tracing in humans

**DOI:** 10.64898/2026.02.12.705660

**Authors:** Teng Gao, Chen Weng, Isaac Johnson, Michael Poeschla, Jonas Gudera, Emily King, Christopher Rouya, Adriana Donovan, Lauren Bourke, Ying Shao, Eladio Marquez, Rahul Tyagi, Leonard I. Zon, Jonathan S. Weissman, Vijay G. Sankaran

**Affiliations:** Division of Hematology/Oncology, Boston Children’s Hospital, Harvard Medical School, Boston, MA, USA; Department of Pediatric Oncology, Dana-Farber Cancer Institute, Harvard Medical School, Boston, MA, USA; Howard Hughes Medical Institute, Boston, MA, USA; Broad Institute of MIT and Harvard, Boston, MA, USA; Whitehead Institute for Biomedical Research, Cambridge, MA, USA; Department of Biology, Massachusetts Institute of Technology, Cambridge, MA, USA; Harvard Stem Cell Institute, Cambridge, MA, USA; Branch Biosciences, Cambridge, MA, USA

## Abstract

Somatic mutations in mitochondrial DNA (mtDNA) provide natural barcodes that enable engineering-free lineage tracing in human tissues, but the complex dynamics of mtDNA inheritance across cell divisions and incomplete sampling of mtDNA introduce uncertainty in reconstructed lineages. Here, we present MitoDrift, a probabilistic framework that integrates Wright–Fisher drift dynamics with sparse single-cell measurements to produce confidence-refined lineage trees enriched for accurate clonal relationships. Validation with gold-standard lentiviral barcoding and whole-genome sequencing demonstrates that MitoDrift outperforms existing tree reconstruction methods in precision while maintaining high clonal recovery, enabling robust analyses linking lineage to cell state. Applying MitoDrift to human hematopoiesis reveals an age-associated decline in clonal diversity with differential impact across cell types and identifies heritable regulatory programs in hematopoietic stem cells *in vivo*, linking AP-1/stress-associated programs to clonal expansions. In multiple myeloma, MitoDrift captures therapy-associated clonal remodeling undetectable by copy number analysis, revealing phenotypic transitions and linking gene regulatory programs to differential drug sensitivity. Collectively, MitoDrift enables high-precision lineage tracing at scale and establishes quantitative lineage–state analysis in primary human tissues, linking clonal history to transcriptional and epigenetic programs in tissue homeostasis, aging, and disease.

## Introduction

Through the lens of descent, evolutionary biology uses ancestry as a scaffold to explain how phenotypes arise and change. The same principle holds in somatic tissues, which are built and maintained through cellular lineages. Somatic mutations that naturally accumulate in a cell’s genome serve as endogenous clonal barcodes, enabling retrospective lineage tracing directly in primary human tissues. Compared to the nuclear genome, mitochondrial DNA (mtDNA) is compact and somatically mutates at a higher rate (>10-fold), making it a highly informative lineage recorder that is scalable and readily integrated with single-cell measurements of cell state^1–3^. Recent advances in mtDNA-based lineage tracing have provided critical insights into human biology, linking clonal identity to functional heterogeneity across homeostasis, aging, and cancer^3–5^. However, quantitative analyses connecting lineage with cell state have yet to be achieved with mtDNA-based lineage tracing, limiting our ability to decipher key processes such as clonal dynamics, heritable gene programs, and cell-state transitions that underlie tissue homeostasis, aging, and disease^6–8^.

Fundamental features of mitochondrial biology pose critical challenges for accurate lineage reconstruction. Mitochondrial genomes are multicopy, turn over actively, and stochastically segregate during division, allowing heteroplasmic mutations to drift toward fixation or become lost in descendants^1,2,9^. Recent methods have advanced mtDNA-based phylogenetic inference but do not account for mitochondrial inheritance dynamics and incomplete sampling^10,11^. Addressing these challenges requires new frameworks to model how mtDNA mutations arise and drift across cell division history. We reasoned that approaches from population genetics could be adapted to the problem of mtDNA-based single-cell lineage inference^12,13^.

Here, we present MitoDrift, a probabilistic framework that treats mitochondrial lineage tracing as an intracellular population-genetic process observed through sparse single-cell measurements. MitoDrift integrates a Wright–Fisher model of heteroplasmy drift with observation noise and performs likelihood-based inference over tree spaces to quantify posterior clade support, yielding confidence-refined cell lineage trees. Benchmarking against orthogonal ground truth from lentiviral barcoding and whole-genome sequencing shows that MitoDrift achieves high-precision lineage inference while maintaining high clonal recovery, outperforming existing methods. In human hematopoiesis, we quantify age-associated oligoclonality across blood lineages and identify heritable regulatory programs in hematopoietic stem cells (HSCs) linked to clonal expansion. In cancer evolution, we resolve treatment-associated clonal remodeling in multiple myeloma (MM) and use phylogeny–state analysis to distinguish heritable from plastic tumor programs, nominating cell states associated with therapy resistance. MitoDrift establishes a scalable platform for quantitative lineage–state analysis linking cell ancestry to transcriptional and chromatin-state programs, yielding new insights into human biology across health and disease.

## Results

### Integrating allelic drift models with single-cell phylogenetic inference

Somatic mtDNA variants provide endogenous barcodes of clonal history, but the resulting lineage signals can be confounded by heteroplasmy dynamics and measurement noise^1,2^. Specifically, heteroplasmic variants can drift toward fixation or extinction across cell divisions, while sparse sampling, contamination, and technical artifacts introduce dropouts and false positives in variant detection (**Figure 1A**). We model this joint process as a hidden Markov tree (HMT): heteroplasmy evolves along lineage paths via Wright–Fisher drift, and latent heteroplasmy states are linked to observed per-cell variant allele counts through a likelihood that jointly captures mutagenesis, drift, and detection error. Given this generative model, MitoDrift computes the tree likelihood via message passing, learns drift and error parameters by expectation–maximization, and quantifies branch uncertainty by Markov chain Monte Carlo (MCMC) sampling over tree space (**Figure 1B**)^14,15^. This procedure produces an initial Neighbor-Joining phylogeny with posterior support (confidence) for each branch; collapsing low-confidence branches yields a refined tree topology enriched for accurate clades, hereafter termed confidence-refined tree (**Methods**; **Figure 1C**). The resulting high-precision lineage trees can be used for quantitative analyses of lineage and cell state, including measurements of clonal diversity, heritability of regulatory modules, and cell state dynamics by linking lineage history with multiomic cell states (**Figure 1D**).

**Figure 1.**
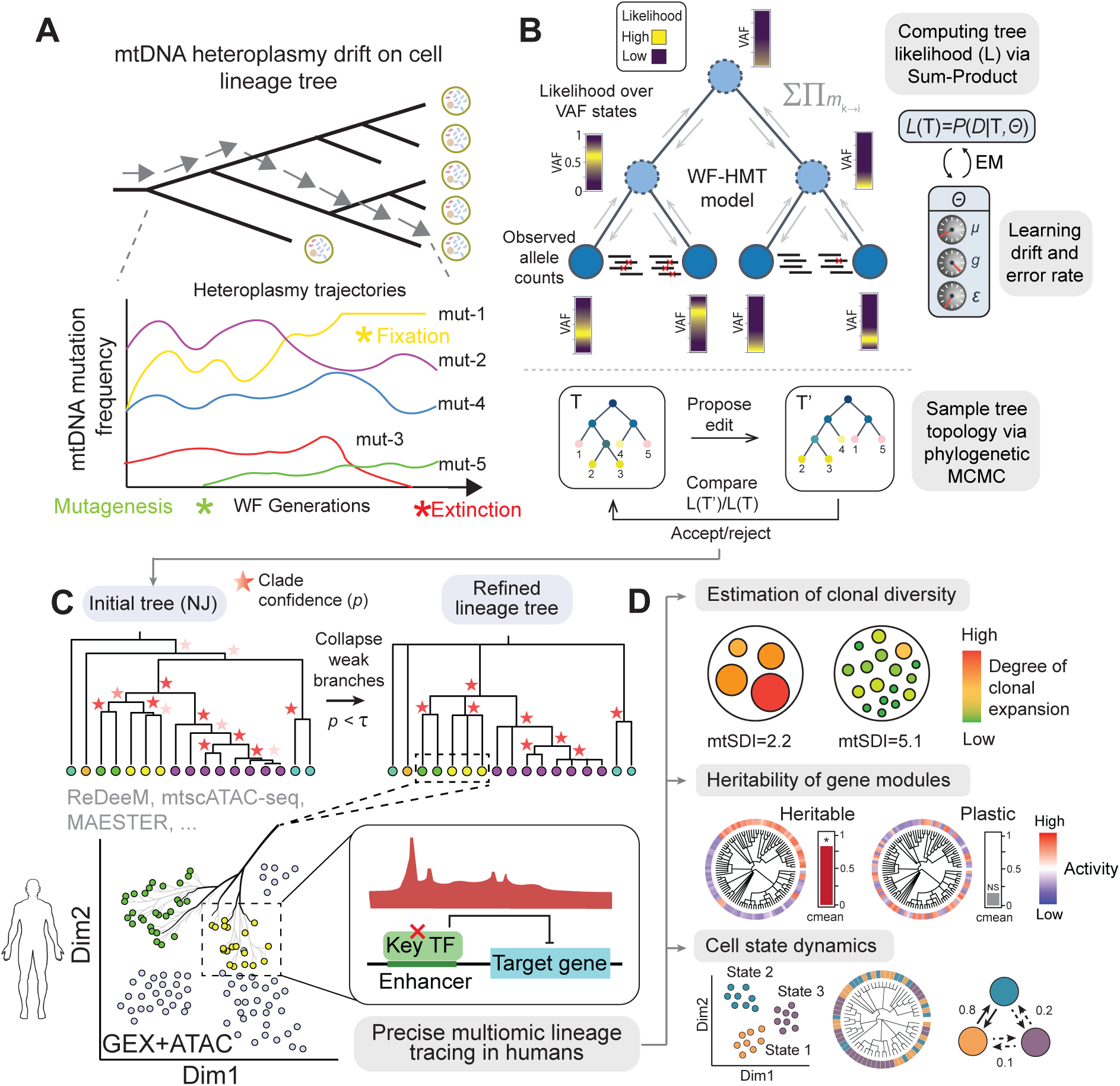
Algorithm overview. (A) Conceptual schematic of mtDNA heteroplasmy dynamics along a cell lineage tree: new mutations arise, heteroplasmy drifts via Wright–Fisher sampling over cell divisions, and lineages experience fixation or extinction of alleles. (B) Probabilistic formulation and inference workflow. For each locus, latent heteroplasmy states at internal nodes are modeled on a discrete VAF grid using a Wright–Fisher hidden Markov tree (WF-HMT), with an observation model for noisy allele counts; tree likelihood is computed by sum–product message passing, model parameters (drift and error rates) are learned by EM, and tree topology is explored using phylogenetic MCMC with accept/reject steps based on likelihood ratios. (C) Confidence-based tree topology refinement and integration with multiome readouts: posterior clade confidence is used to collapse weakly supported branches in an initial tree (e.g., Neighbor-Joining) to produce a refined lineage tree at a chosen cutoff τ, enabling joint analysis of lineage hierarchies and multiome regulatory features. (D) MitoDrift enables several *in vivo* analyses of lineage and cell state in human tissues, including estimating clonal diversity/expansion (e.g., Shannon diversity index, SDI), quantifying heritability versus plasticity of gene modules/regulons, and inferring cell-state couplings and transitions by connecting lineage history with multiomic cell states.

### Lentiviral barcoding validates high-fidelity clonal reconstructions

To benchmark MitoDrift for mtDNA-based lineage inference, we applied lentiviral barcoding (lineage and RNA recovery, LARRY v2)^16^ to primary human HSCs prior to expansion and differentiation, then performed single-cell regulatory multiomics with deep mtDNA profiling (ReDeeM) on the expanded progeny with simultaneous readout of LARRY barcodes. This approach provided matched measurements of exogenous barcode-defined clones and endogenous mtDNA variants in each cell, enabling direct evaluation of mtDNA lineage inference against ground-truth clonal relationships (**Figure 2A**; **Methods**). We inferred lineage trees from mtDNA variants using MitoDrift and evaluated concordance with ground-truth clones defined by lentiviral LARRY barcodes (hereafter termed Lenti-clones) using a clade-level precision–recall framework (**Methods**). We treat each internal node in the inferred tree as a predicted clade and evaluate its accuracy by the Jaccard similarity index (J) between a clade’s tip set and Lenti-clones. We define clade precision as the fraction of inferred clades that match a Lenti-clone with Jaccard similarity exceeding a specified threshold, and clone recall as the fraction of Lenti-clones that are recovered by a predicted clade exceeding the same Jaccard threshold. This framework captures the tradeoff between resolving fine-grained structure (higher recall) and avoiding spurious splits (higher precision) when inferring lineage trees.

**Figure 2.**
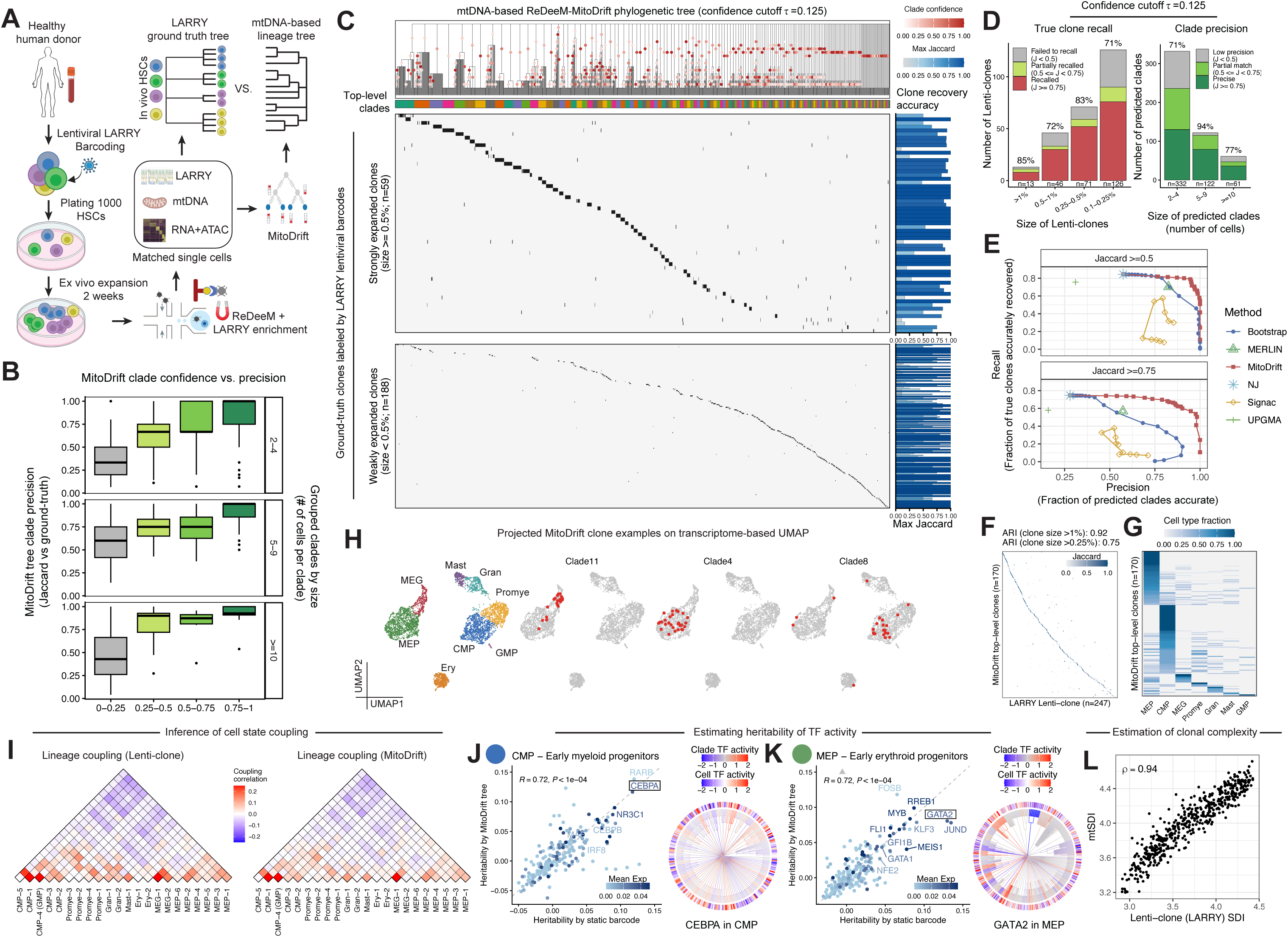
Benchmarking MitoDrift using lentiviral barcoding and validating lineage-state analyses. (A) Experimental and computational overview for benchmarking: human HSCs are lentivirally barcoded (LARRY), profiled by scMultiome with deep mtDNA capture (ReDeeM), and matched single cells are used to infer an mtDNA-based MitoDrift lineage tree for comparison to the static-barcode ground-truth tree. (B) Relationship between MitoDrift clade confidence and clade precision across clade size bins (clades grouped by number of cells). (C) Example MitoDrift phylogeny (refined at clade confidence threshold τ = 0.125; singletons omitted) annotated with inferred clones (top-level clades) and a cell-by-clone agreement view against LARRY Lenti-clone ground truth. (D) True clone recall and MitoDrift clade precision summarized across clone-size strata using Jaccard similarity between inferred clades and ground-truth clones using the refined tree at τ = 0.125. J, Jaccard similarity. (E) Precision–recall performance of lineage inference methods on subsampled datasets (45 clones per replicate, 10 replicates) evaluated using different clade matching stringency (Jaccard≥0.5 and Jaccard≥0.75); for MitoDrift, curves are generated by sweeping τ. (F) Clone-to-clone overlap between MitoDrift top-level clades (τ = 0.125; singletons omitted) and LARRY clones, colored by overlap fraction (Jaccard); adjusted Rand index (ARI) at different clone size cutoffs. (G) Cell-type fraction per MitoDrift top-level clones (τ = 0.125; singletons omitted). (H) Projection of representative MitoDrift top-level clades onto transcriptome-based UMAP, illustrating clade distribution across hematopoietic states. (I) Lineage coupling between cell differentiation states inferred from static barcodes versus MitoDrift trees under a Brownian-motion distance model. (J) Validation of TF-activity heritability estimates in CMP by comparing heritability computed from LARRY ground-truth lineages and MitoDrift lineages, with CEBPA shown as an example TF (tip TF activity and clade-level TF z-score mapped to the lineage tree; singletons omitted). (K) Same analysis in MEP, with GATA2 shown as an example TF. (L) Comparison of clonal diversity estimates using static barcodes and MitoDrift phylogenies on subsampled datasets (∼300 cells per replicate, 200 replicates with clone-level subsampling).

We first asked whether clade confidence estimated by MitoDrift provides a means to calibrate lineage inference resolution and accuracy. Across clade sizes, clades with higher confidence showed higher empirical precision when evaluated against Lenti-clones (**Figure 2B**). Leveraging this relationship, collapsing low-confidence branches in the initial tree topology by threshold т = 0.125 yields a tree structure that accurately aligns with barcode ground-truth (**Figure 2C**; **Figure S1A-C, Figure S2**; **Methods**). The confidence-refined tree accurately recovers 75% of Lenti-clones (J ≥ 0.5), with consistent performance across Lenti-clone sizes: 85% recall for strongly expanded clones (each comprising ≥1% of total cells, n=13), 79% for moderately expanded clones (0.25–1%, n=117), and 71% for weakly expanded clones (0.1–0.25%, n=126; **Figure 2D**; **Figure S1**). Importantly, the refined tree achieved 77% overall clade precision, substantially outperforming traditional phylogenetic methods (Neighbor-Joining, 55%; UPGMA, 28%; **Figure 2E**; **Figure S1**). Further increasing the confidence threshold achieves even higher clade precision (90% precision at 70% clone recall, т = 0.3; **Figure S1D,E**). Across confidence thresholds, MitoDrift demonstrated superior precision–recall performance compared to existing tree inference methods (**Figure 2E**). Clonal partitions from the MitoDrift phylogeny demonstrate high concordance with Lenti-clones (clone size >1%: adjusted Rand index [ARI] = 0.92; clone size >0.25%: ARI = 0.75; **Figure 2F**; **Figure S1F**; **Methods**).

### MitoDrift enables robust lineage–state analysis

We then sought to evaluate the capability of lineage–state analyses using MitoDrift phylogenies, validating against the Lenti-clone ground truth measurements. We focused on three key aspects of such analysis: (1) clonal fate bias and cell-state coupling, (2) heritability of gene regulatory programs, and (3) clonal population structure and diversity.

First, we observe that most top-level clades in the MitoDrift phylogeny are strongly enriched for specific hematopoietic cell types (**Figure 2G,H**), revealing pronounced clonal fate bias in human HSCs and multipotent progenitor cells (MPPs) *in vitro*, echoing mouse HSPC barcoding studies^17,18^. This observation prompted us to ask whether MitoDrift phylogenies can resolve cell-state coupling along the differentiation hierarchy. Using MitoDrift phylogenies with PATH^19^ to infer lineage coupling between fine-resolution differentiation stages, we recovered coupling structure highly concordant with estimates derived from barcode ground truth (R = 0.96, CCC = 0.91; **Figure 2I, Figure S3B**). MitoDrift’s precise tree topology improved both the accuracy and robustness of cell-state coupling estimation compared to Neighbor-Joining (**Figure S3B**).

Second, determining which phenotypic traits are heritable across cell divisions and which traits are transient or plastic is a long-standing question in stem cell biology and cancer evolution. To assess whether mtDNA-based lineage trees can identify heritable regulators of lineage specification, we combined MitoDrift phylogenies with gene regulatory network inference SCENIC+^20^ to estimate heritability of transcription factor (TF) activity across the differentiation landscape. Regulon heritability estimates derived from MitoDrift trees closely matched those computed from barcode ground truth measurements in common myeloid progenitors (CMPs; R = 0.72, p < 1e-4) and megakaryocyte-erythroid progenitors (MEPs; R = 0.72, p < 1e-4; **Figure 2J,K**). This analysis highlighted cell-type-specific regulators such as *CEBPA* in CMPs and *GATA2* in MEPs with pronounced clonal heritability in regulon activity (**Figure 2J,K**). To further explore whether heritable regulatory variation at the progenitor stage prefigures downstream fate, we reanalyzed an earlier mtscATAC-seq dataset^2^, which profiles *in vitro* CD34+ HSPC differentiation across three timepoints (days 8, 14, and 20; *n* = 10,414 cells; **Figure S4A,B**). MitoDrift inferred a phylogeny that, when integrated with chromatin-derived cell states using PATH, revealed the expected myeloid–erythroid bifurcation (**Figure S4C**). We then focused on day 8 progenitors (n = 656), reasoning that heterogeneity at this early stage may prefigure downstream fate. We first used MitoDrift phylogenies to test whether progenitor regulatory states exhibit phylogenetic signal indicative of lineage priming. Both myeloid and erythroid pseudotime scores—derived from day 8 chromatin accessibility—showed significant phylogenetic signal, quantified by cmean (myeloid: cmean = 0.22, P < 2 × 10^−4^; erythroid: cmean = 0.23, P < 2 × 10^−4^). Clades within the day 8 phylogeny showed distinct enrichment for myeloid- versus erythroid-primed states (**Figure S4D,E**). Phylogenetic signal analysis of chromVAR-derived TF activities identified AP-1/bZIP, C/EBP (myeloid differentiation), GATA (erythroid specification), and stem cell–associated modules (HLF, BACH) as exhibiting the strongest heritability (**Figure S4F**). We next asked whether progenitor state predicts downstream fate. We quantified lineage bias by measuring each day 8 progenitor’s phylogenetic proximity to myeloid versus erythroid cells at days 14 and 20 (**Methods**). Using this approach, we found that myeloid pseudotime at day 8 was predictive of subsequent myeloid bias among descendant cells (**Figure S4G**). Consistent with known lineage regulators, regression analysis identified C/EBP family members and *SPI1* (PU.1) as positively associated with myeloid output, whereas GATA factors were associated with erythroid fate (**Figure S4H**). Together, these analyses demonstrate that MitoDrift’s confidence-refined phylogenies can connect early progenitor regulatory states to downstream differentiation outcomes.

Finally, to benchmark the accuracy of clonal diversity estimation using MitoDrift phylogenies, we generated simulated datasets spanning a range of clonal diversities by subsampling different sets of Lenti-clones (**Methods**). We then computed the mtDNA-based Shannon diversity index (mtSDI) from MitoDrift trees and found strong agreement with barcode-derived estimates (ρ = 0.94; **Figure 2L; Methods**). These results demonstrate that MitoDrift can recover clonal history with high precision, establishing quantitative phylogeny–state analysis with single-cell multiome data.

### MitoDrift resolves clonal dynamics in native hematopoiesis

To benchmark MitoDrift lineage inference in the context of native human hematopoiesis, we employed a whole-genome sequencing (WGS) dataset of single colonies derived from HSCs/MPPs in eight healthy donors spanning 29–81 years of age (**Figure 3A**)^7^. In this setting, WGS-based phylogenies reconstructed from nuclear SNVs provide a time-resolved orthogonal ground truth for evaluating mtDNA-based lineage trees (**Figure S5**; **Methods**). We focused on clades representing post-natal clonal expansions, defined as those occurring after 100 mutations of molecular time^7^ (**Figure S5C,D**). Starting from the initial Neighbor-Joining tree (21% recall at 6% precision; J ≥ 0.5), MitoDrift’s confidence-based topology refinement significantly improved clade precision while maintaining recall, recovering ∼13% of clades with 50% precision and ∼10% of clades with 70% precision. Across the full precision–recall curve, MitoDrift showed the most favorable performance, exceeding MERLIN and bootstrap-based approaches (**Figure 3B**). We note that *in vitro* colony expansion could lead to loss of mtDNA variants, and that conventional WGS has limited sensitivity for detecting low-VAF mtDNA heteroplasmy — both factors make this benchmark a lower bound on recoverable lineage signal^4^.

**Figure 3.**
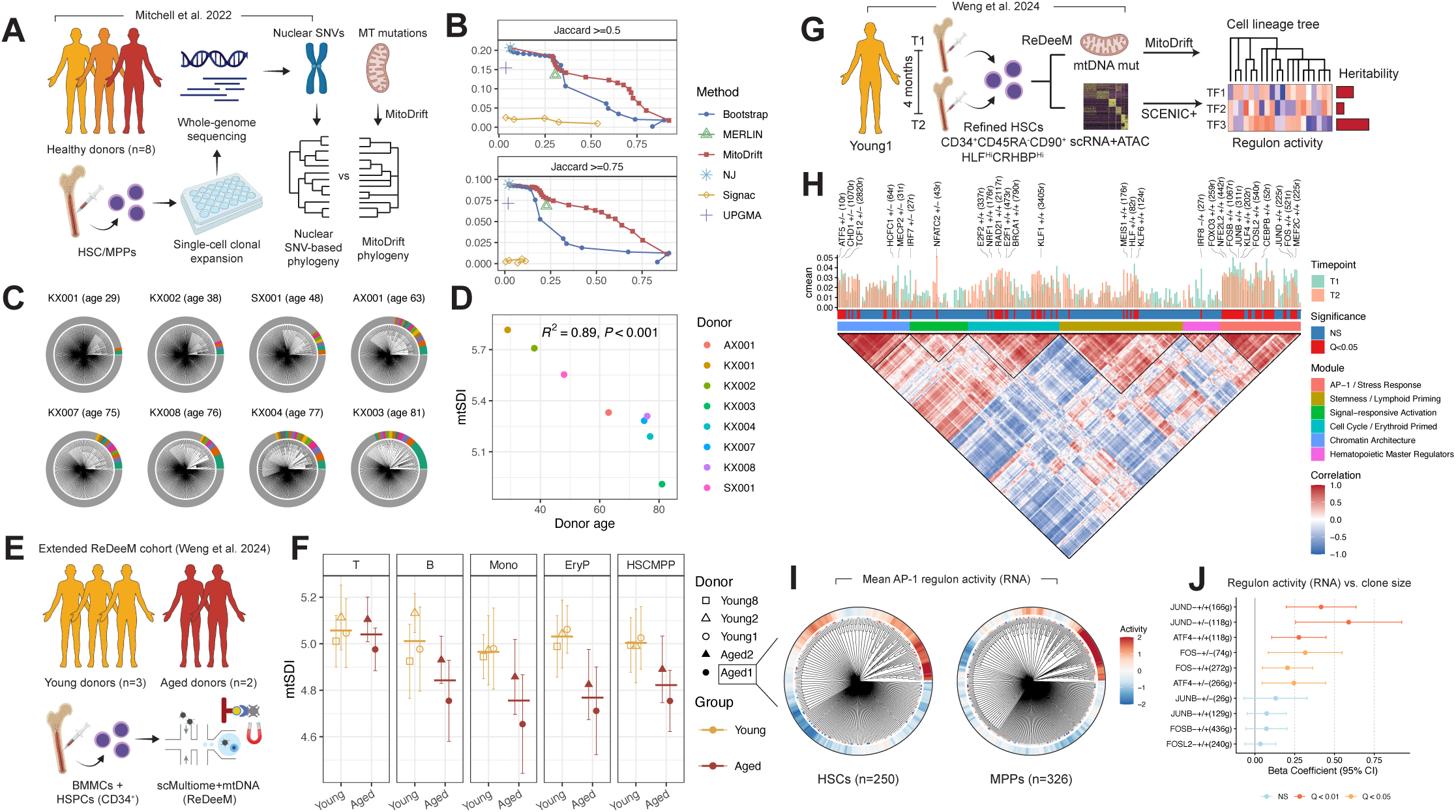
Validation of MitoDrift *in vivo* and application to uncover clonal memory of HSC regulatory programs in steady-state hematopoiesis. (A) Benchmarking design: mtDNA mutations are used to infer lineage trees of donor HSCs/MPPs profiled by whole-genome sequencing of single-cell clonal expansions, and accuracy is evaluated against nuclear SNV-based phylogenies (derived from WGS data). (B) Precision–recall curves for lineage inference methods on subsampled datasets (150 cells per donor per replicate, 10 replicates) evaluated at Jaccard ≥ 0.75 and Jaccard ≥ 0.5; for MitoDrift, curves are generated by sweeping τ. (C) Per-donor MitoDrift phylogenies for the WGS benchmark cohort (healthy donors, n = 8; refined at confidence threshold τ = 0.05), shown across donors with ages indicated at the top. The outer ring highlights top-level clades representing >1% of cells. (D) Association between mtDNA-based clonal complexity (mtSDI) and donor age. (E) Extended ReDeeM cohort: study design for scMultiome+mtDNA profiling of young (n = 3) and aged (n = 2) donors from BMMCs and CD34+ HSPCs. (F) mtSDI in major mature blood cell and progenitor compartments, stratified by donor and age group; computed from 20 random subsamples of 200 cells per cell type per donor (mean ± 95% CI). (G) Longitudinal multiome+mtDNA lineage analysis schematic for refined HSCs, combining MitoDrift lineage inference with SCENIC+ regulon activity to quantify heritability of HSC state regulation. (H) Heritability of HSC regulons across timepoints measured by ATAC signal. From top to bottom: heritability (cmean) of regulons at timepoint T1 and T2; statistical significance (NS versus Q < 0.05); functional module assignments; correlation of regulon activity across cells. (I) Mean AP-1 regulon activity visualized for Aged 1 HSCs and MPPs. (J) Regression of AP-1–associated regulon activity against clone size in donor Aged 1. Effect sizes are reported as beta coefficients with 95% CIs. Colors indicate significance levels based on Q values.

We then asked whether MitoDrift phylogenies can detect age-associated changes in HSC clonal diversity. Across 8 donors in the WGS benchmark cohort, mtDNA-based clonal diversity declined significantly with donor age, consistent with WGS-based measurements (R = 0.89, P < 0.001; **Figure 3C,D**). To examine how clonal diversity changes in HSCs impact different hematopoietic cell types, we utilized the ReDeeM dataset from Weng et al.^4^ with one additional donor to examine mtSDI across hematopoietic cell types as annotated by the corresponding multiome data (**Figure 3E**). This cohort comprised young (n = 3) and aged (n = 2) donors profiled from bone marrow mononuclear cells (BMMCs) and CD34+ hematopoietic stem and progenitor cells (HSPCs). As expected, older donors showed reduced mtSDI in the HSC/MPP compartment compared to younger donors, consistent with the single-colony WGS results (**Figure 3F**). Notably, mtSDI of T cells did not exhibit a pronounced age-related decrease, whereas B cells, monocytes, and erythroid progenitors showed marked reductions in mtSDI with age (**Figure 3F**). We further confirmed these cell-type-specific clonal diversity patterns by applying MitoDrift to an additional multi-donor ReDeeM dataset and a clonal hematopoiesis sample profiled by enrichment of mtDNA-encoded transcripts in single-cell RNA-sequencing data^3,4^ (MAESTER; **Figure S6A-C**). This observation is consistent with recent nuclear-mutation lineage tracing showing that expanded HSC clones in aged humans often exhibit myeloid-biased or myeloid/B-biased output with reduced T cell output^21,22^; the relative preservation of T cell clonal diversity may reflect the long-lived nature of memory T cells in the bone marrow and their reduced dependence on ongoing HSC contributions (**Figure S6D**). Taken together, MitoDrift enables quantitative studies of clonal dynamics in native human hematopoiesis, revealing its decline in HSC aging and demonstrating differential impacts across hematopoietic cell types.

### Mapping heritable HSC regulatory programs and signatures of clonal expansion

Next, we assessed whether MitoDrift phylogenies can reveal heritable gene regulatory programs in HSCs *in vivo*—a fundamental question for understanding clonal selection and functional heterogeneity. We reanalyzed serial ReDeeM profiling data of refined HSCs from a healthy young donor^4^ and integrated MitoDrift lineage inference with SCENIC+^20^ to estimate regulon heritability using RNA and ATAC-derived activity scores (**Figure 3G; Figure S7; Methods**). Significant regulon heritability was detected using ATAC-derived chromatin accessibility scores, but not RNA-based expression levels (**Figure S8**). Based on covariation across cells, ATAC-based regulon activity organized into functional modules including AP-1/stress response, stemness/lymphoid priming, signal-responsive activation, cell cycle/erythroid priming, chromatin architecture, and hematopoietic master regulators (**Figure 3H**). Heritable regulons (Q < 0.05) were distributed across multiple modules, including AP-1/stress response (*FOS*, *FOSB*, *FOSL2*, *JUNB*, *JUND*, *NFE2L2*, *FOXO3*, *KLF4*, *CEBPB*), stemness/lymphoid priming (*HLF*, *MEIS1*, *KLF6*), erythroid-primed cycling (*E2F1*, *E2F2*, *BRCA1*, *RAD21*, *NRF1*, *KLF1*), and chromatin-organization programs (*CHD1*, *MECP2*, *HCFC1*, *TCF12*) (**Figure 3H**). Together, these results support the presence of stable, lineage-associated regulatory programs *in vivo* spanning stress response, lineage priming, and chromatin regulation. We also note that detection power due to sparse clonal sampling and limited lineage resolution may contribute to lower or less consistent heritability estimates for some regulons.

Although age-related clonal expansion of HSCs has recently been shown by WGS-based lineage tracing using somatic mutations^7,23,24^, it remains unclear whether clonally expanded HSCs are phenotypically distinct and what mechanisms may drive clonal dominance. Inflammatory pathways, particularly the AP-1 family of transcription factors, have been implicated in HSC aging^25–27^, but their relationship to clonal expansion remains unclear. To investigate this, we used MitoDrift to reconstruct clonal structures in HSCs and MPPs from two aged donors in the ReDeeM cohort, and assessed their association with AP-1 regulon activity (**Methods**; **Figure 3F**). Interestingly, RNA-based transcriptional activity of AP-1 factors was significantly associated with clone size in HSCs/MPPs of Aged 1 (Q = 4.5×10⁻⁴; regression beta = 0.24; **Figure 3I,J**; **Figure S9**). Together, these analyses demonstrate the use of MitoDrift for quantitative lineage-state analysis in native hematopoiesis, revealing heritable gene regulatory programs in human HSCs at steady-state and identifying the top heritable program, AP-1, as being associated with age-related clonal expansions.

### Tracking treatment-associated clonal remodeling in multiple myeloma

Cancer therapy resistance and disease relapse remain among the most critical challenges in oncology, yet the clonal dynamics and evolutionary paths of surviving cells that repopulate tumors after treatment remain unclear. Multiple myeloma is a plasma cell malignancy that nearly always relapses after initial response and can be longitudinally sampled over extended treatment courses, making it an ideal disease context to study the roles of clonal remodeling and cell-state plasticity in therapy resistance^28,29^. While copy number alterations (CNAs) define major genetic subclones, the fine-grained clonal structure within CNA-defined populations and its relationship to divergent tumor cell states and drug sensitivity remain poorly understood^28–30^. Previous efforts using mtDNA mutations to track tumor evolution provide valuable insights, yet are limited in resolution^5,28^. We therefore applied MitoDrift to a longitudinal ReDeeM dataset, composed of three multiple myeloma patients sampled at diagnosis and following induction therapy (**Figure 4A**). Across patients (MM1-3), post-treatment tumor burden (plasma cell percentage) decreased to varying extents—with the largest reduction in MM1, intermediate in MM2, and minimal in MM3 (**Figure 4B**). To characterize the cellular landscape across timepoints, we performed weighted nearest-neighbor (WNN) integration of RNA and ATAC modalities and projected cells onto a unified UMAP embedding. This analysis revealed major hematopoietic populations including T cells, B cells, NK cells, monocytes, and plasma cells (**Figure 4C**). Malignant plasma cells were identified based on CNAs inferred from the multiome data (**Figure S10A**; **Methods**).

**Figure 4.**
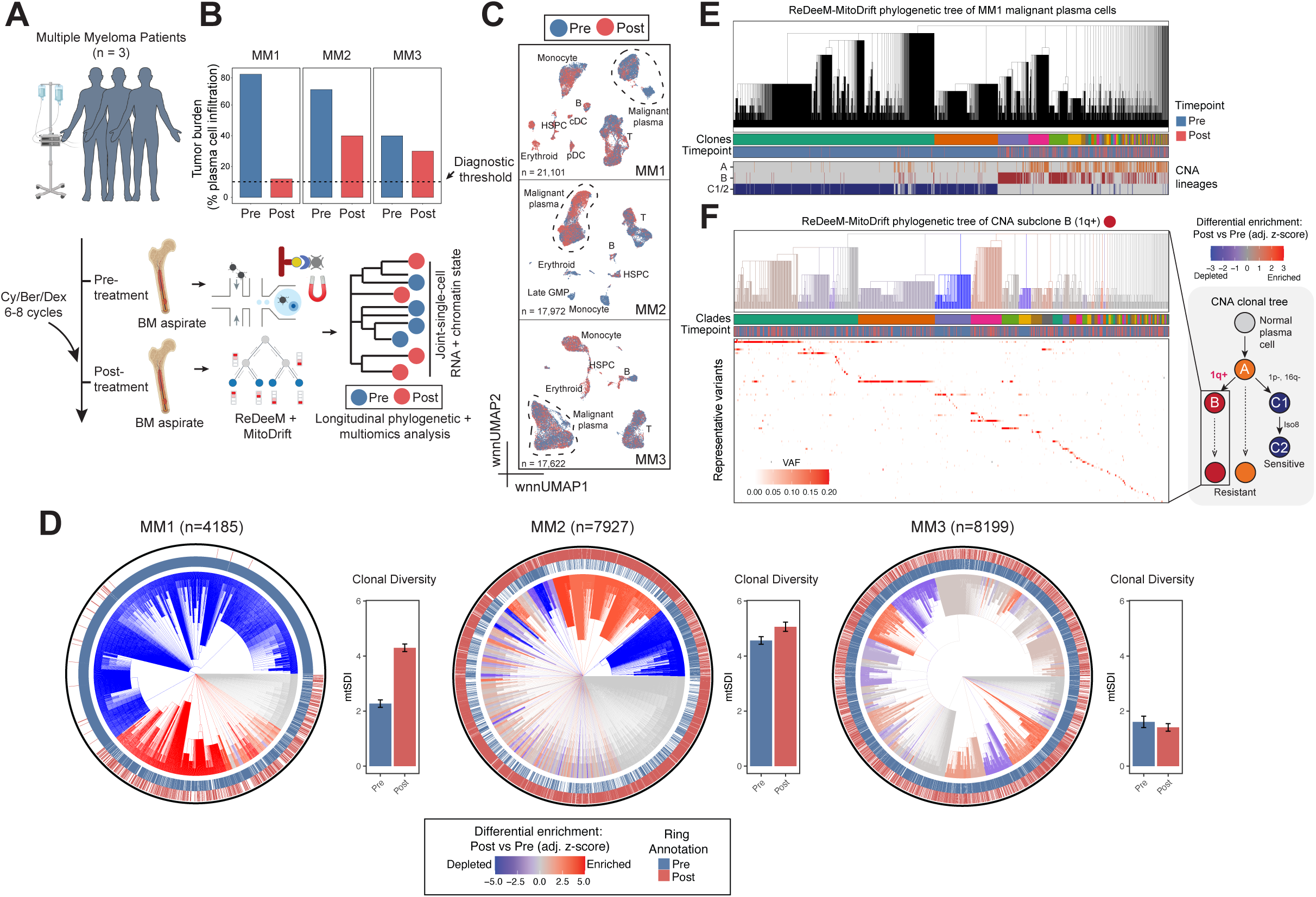
Longitudinal phylogenetic multiome profiling in multiple myeloma captures clonal remodeling under therapy. (A) Study design and cohort overview for longitudinal sampling of multiple myeloma patients (pre- and post-treatment), including ReDeeM multiome+mtDNA profiling and MitoDrift lineage inference. (B) Tumor burden (% plasma infiltration) by donor and timepoint, with a diagnostic threshold indicated. (C) wnnUMAP (ATAC+RNA) projections across donors showing major cell compartments colored by timepoint (Pre vs Post). The encircled cells contain mostly malignant plasma cells and a small number of normal plasma cells, identified based on inferred CNAs. (D) MitoDrift-inferred phylogenies of malignant plasma cells per donor, with branches colored by differential enrichment for post- versus pre-treatment cells and ring annotations indicating sampling timepoint; clonal diversity summaries compare pre- versus post-treatment malignant plasma cells. (E) ReDeeM–MitoDrift phylogeny of donor MM1 malignant plasma cells (refined at confidence threshold τ = 0.075; singletons omitted) annotated with sampling timepoint and CNA-defined lineages. For visualization, CNA subclones C1 and C2 (C2 derived from C1) are grouped as a single CNA lineage (C1/C2). (F) MitoDrift-inferred phylogeny within the 1q+ CNA subclone (subclone B) showing fine-scale clade structure and timepoint-associated enrichment.

We next applied MitoDrift to reconstruct lineage trees of malignant plasma cells within each patient. Across patients, the inferred phylogeny revealed hierarchical clonal structures with differential enrichment for post- versus pre-treatment cells, suggesting therapy-related clonal remodeling and lineage-specific drug sensitivity (**Figure 4D**; **Figure S11**). To assess concordance between MitoDrift phylogenies and CNA-defined clonal structure, we compared inferred clades to CNA-defined subclones in the tumor of MM1, which harbors pronounced CNA subclonal heterogeneity (**Figure 4E; Figure S10B,C**). CNA-defined subclonal lineages appeared as continuous blocks on the mtDNA phylogeny, with high phylogenetic autocorrelation (Moran’s I = 0.41-0.76), indicating that mtDNA phylogenies capture major axes of genetic divergence defined by CNAs (**Figure 4E**). To test whether mtDNA lineages can enable finer-grained analysis of evolution within CNA-defined tumor subpopulations, we focused on the therapy-resistant subclone harboring 1q-gain, a recurrent CNA in MM associated with adverse prognosis^31^ (**Figure 4F**). Within this subclone, MitoDrift revealed further clonal diversification not captured by CNA analysis. These subclades showed distinct temporal distributions, suggesting subclonal heterogeneity in drug sensitivity. Notably, in patients MM2 and MM3, which lacked pronounced CNA subclonal heterogeneity, MitoDrift nonetheless resolved clades with distinct temporal distributions, suggesting that MitoDrift lineages can detect therapy-related clonal remodeling even in the absence of detectable CNA subclones (**Figure 4D**).

Using the MitoDrift tumor phylogenies, we quantified clonal diversity at each timepoint (**Figure 4D**; **Figure S11**). Notably, only patient MM1, the deep responder, showed a marked increase in tumor clonal diversity after therapy (mtSDI = 2.3, pre-treatment; mtSDI = 4.3, post-treatment), whereas patients MM2 and MM3 maintained similar tumor clonal diversities post-treatment. The elevated post-treatment clonal diversity in MM1 is consistent with the eradication of dominant clones by therapy (**Figure 4E**) but also suggests that the persistent disease came from a polyclonal reservoir rather than a single expanded clone, as directly observed in the subclonal phylogeny (**Figure 4F)**. Together, these results demonstrate that MitoDrift can resolve fine-grained clonal structure and therapy-relevant subclonal heterogeneity that extends beyond the resolution of CNA analysis, and highlight the potential of mtDNA-based phylogenies to identify patients with distinct evolutionary trajectories.

### Linking heritable and plastic tumor states to therapy resistance

Linking gene programs to clonal structure and tracking their dynamics during therapy is key to understanding resistance mechanisms. Here, we integrated MitoDrift phylogenies with multiomic data to perform phylogeny–state analysis in malignant plasma cells, quantifying cell-state heritability and transition dynamics before and after treatment (**Figure 5**). We first characterized the cell-state landscape of malignant plasma cells in donor MM1, using WNN integration of RNA and ATAC modalities to define cell states (**Figure 5A**; **Methods**). Clustering identified nine cell states spanning a range of functional programs, including dormant/signaling-dampened (P1), normal-like plasma (P2), adaptive signaling/stress-buffered (P3), inflammatory chemokine-secreting (P4), lipid-inflammatory remodeling (P5), cytokine-secreting (P6), proliferative (P7), dedifferentiated stress-tolerant (R1), adaptive metabolic-adhesive (R2), and CD34+ progenitor-like (R3) phenotypes (**Figure 5A**). We then examined major clades (A–F) in the MitoDrift phylogeny and examined their distributions across tumor cell states and treatment timepoints (**Figure 5B**). Distinct clades show differential enrichment in specific cell states before and after treatment, with some clades enriched in pre-treatment inflammatory cytokine-secreting (P4) states while others connect pre-treatment signaling-dampened (P1) cells with post-treatment dedifferentiated stress-tolerant (R1) cells. Such clade-specific enrichment suggests that lineage history constrains cell-state distributions within certain patients and can be used to infer evolutionary dynamics between tumor cell states.

**Figure 5.**
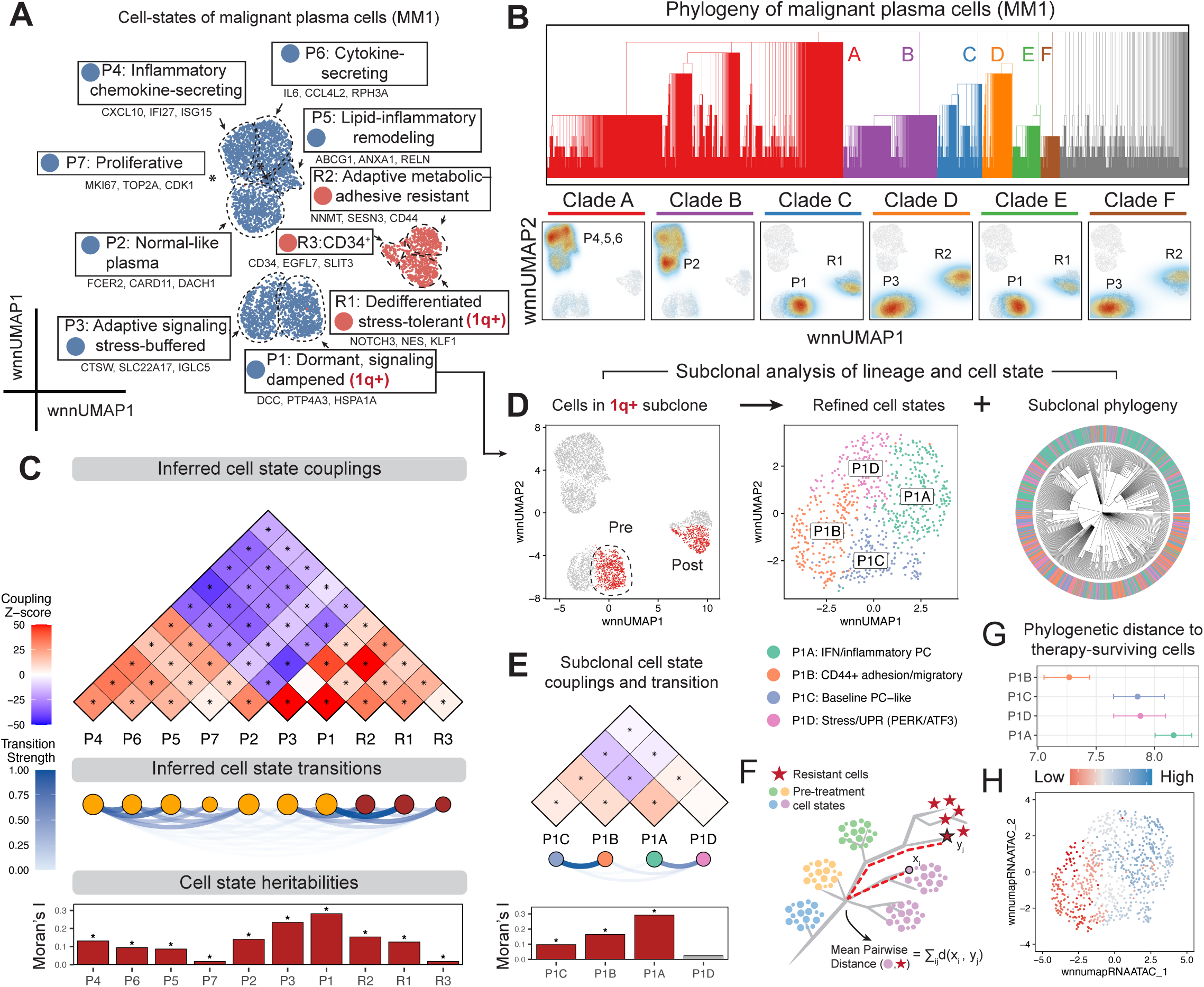
Phylogeny–state analysis links tumor evolution to resistant cell states. (A) Cell state landscape of malignant plasma cells (donor MM1) on a wnnUMAP embedding created using RNA and ATAC measurements. (B) MitoDrift phylogeny of malignant plasma cells of MM1 (refined at confidence threshold τ = 0.075; singletons omitted) partitioned into clades, with clade-specific enrichments across cell states and pre/post treatment. (C) PATH analysis quantifying lineage coupling, cell-state transitions, and cell-state heritability based on the MitoDrift phylogeny and multiome cell states. (D) Subclone-focused analysis combining refined cell states with the subclonal MitoDrift phylogeny (refined at confidence threshold τ = 0.075; singletons omitted). (E) Cell-state couplings and transition dynamics within the 1q+ subclone inferred by PATH using MitoDrift phylogenies. (F) Schematic for quantifying phylogenetic relatedness of pre-treatment cell states to post-treatment cells using inter-community distance. (G) Phylogenetic distance from pre-treatment subclonal cell states to post-treatment cells. (H) Cells within the 1q+ subclone projected onto wnnUMAP and colored by inferred phylogenetic distance to post-treatment cells, highlighting regions associated with higher relatedness (lower distance) to post-treatment cells.

To formally quantify lineage–state relationships, we applied PATH analysis to the MitoDrift phylogeny and multiome cell states in donor MM1 (**Figure 5C**; **Methods**). Examining lineage coupling patterns and transition dynamics between pre-treatment states and post-treatment cells, we found that most pre-treatment states showed weak phylogenetic relatedness to post-treatment cells. However, states P1 and P3 exhibited the strongest coupling and transition strength with post-treatment cells, suggesting these states harbor cells most closely related to the therapy-resistant population. These states are characterized by stress-response and signaling-dampening programs, consistent with phenotypes associated with treatment resistance in other cancer contexts^32,33^. Cell-state heritability varied across states, with cell states P1 and P3 showing the highest heritabilities (Moran’s I = 0.28 and 0.23, respectively; **Figure 5C**). In contrast, pre-treatment state P7 (proliferative) and post-treatment state R3 (progenitor-like) were among the least heritable (Moran’s I = 0.02 for both), indicating that they were plastic or transient cell states. Similar patterns of mixed heritable and plastic cell states with distinct evolutionary relationships to post-treatment cells were also observed in other patients (**Figure S12**). These results demonstrate that MitoDrift phylogenies combined with multiomic readouts can distinguish heritable lineage-restricted programs from plastic variation in malignant plasma cells, and that longitudinal lineage-state analysis can identify gene programs most evolutionarily related to post-treatment cells.

Therapy failure is often driven by rare subclonal "persister" cells that survive treatment through both genetic and non-genetic mechanisms^34,35^. A central challenge is linking subclonal lineage history to cell state to identify resistance programs and track how these programs evolve over time. Here, we focused on the 1q-gain subclone, which showed treatment resistance and subclonal remodeling (**Figure 4F**). However, whether cell-state heterogeneity within this subclone confers differential drug sensitivity remains unclear. Within this subclone, we re-defined cell states using WNN graph-based clustering and applied PATH analysis to the subclonal phylogeny (**Figure 5D,E**). The subclonal cell states showed distinct heritability and transition patterns, with P1A (INF/inflammatory; most heritable: Moran’s I = 0.29) coupled with P1D (Stress/UPR; least heritable: Moran’s I = 0.02) and P1B (CD44+ adhesion/migratory) coupled with P1C (Baseline PC-like). Next, we asked which pre-treatment cell states within the 1q gain subclone are most closely related to post-treatment cells, as these may represent candidate resistance programs. We used inter-community phylogenetic distance as a measure of how different pre-treatment cell states are related to therapy-surviving cells (**Figure 5F**; **Methods**). Among refined pre-treatment states, P1B showed the shortest phylogenetic distance to post-treatment cells, whereas P1A was the most distant (**Figure 5G,H**). The P1B state is characterized by high CD44+ adhesion/migratory signaling, consistent with CD44’s established role in cell adhesion-mediated drug resistance in myeloma, where CD44 engagement with the bone marrow microenvironment confers protection^36,37^. Together, these results demonstrate that MitoDrift phylogenies can link tumor clonal evolution with cell-state dynamics at high resolution, allowing us to nominate resistance-driving gene regulatory programs.

## Discussion

Mitochondrial DNA variants offer a powerful approach for manipulation-free lineage tracing in humans, but the complex dynamics of mitochondrial inheritance have limited confidence and resolution of lineage reconstruction. Existing studies have largely focused on qualitative associations between lineage and cell state by examining coarse clonal contrasts^1–3,5,38^ or descriptive comparisons of clonal complexity across conditions^39^. MitoDrift addresses these limitations by jointly modeling intracellular allelic drift and single-cell measurement noise via a hidden Markov tree model. Events that confound mitochondrial lineage tracing, including mutation loss, recurrence, drift, and false detections, are incorporated into the overall likelihood model when evaluating confidence in the tree topology. Refining the tree topology by collapsing branches with low confidence allows downstream analyses to focus on accurate lineage structures, while acknowledging regions of the tree that remain poorly resolved. Benchmarking against orthogonal ground truths from lentiviral barcoding and whole-genome sequencing demonstrated that MitoDrift enables high-precision lineage inference across biological contexts, establishing the reliability of quantitative lineage–state analysis using mtDNA-based phylogenies.

Applying MitoDrift to human hematopoiesis, we found that mtDNA-based phylogenies capture age-associated oligoclonality with cell-type-specific patterns in steady-state hematopoiesis. T cells maintained clonal diversity with age, whereas myeloid cells, B cells, and erythroid progenitors showed marked reductions. This pattern is consistent with recent observations that dominant HSC clones exhibit myeloid-biased lineage output in aged individuals^21^ and following allogeneic hematopoietic cell transplantation^22^. We note that unlike T-cell receptor diversity in peripheral-blood T cells, which declines with age^40^, mtDNA-based clonal diversity reflects the number of historical hematopoietic ancestors. Because bone marrow T cells are predominantly long-lived memory cells, their clonal signatures preserve historical HSC diversity even as the active progenitor pool contracts. More broadly, when the HSC pool becomes oligoclonal, lineage-biased output from dominant clones can differentially reduce clonal diversity across mature cell types: lineages that receive disproportionate contribution from expanded clones will show reduced diversity, whereas lineages with minimal contribution retain polyclonal ancestry (**Figure S6D**). Reduced clonal complexity in myeloid/erythroid output may reduce hematopoietic redundancy, increasing susceptibility to stress and making peripheral function more dependent on the behavior of a small number of dominant clones. We also identified heritable regulatory programs in HSCs, with AP-1/stress-associated and quiescence-related regulons showing significant phylogenetic autocorrelation across longitudinal sampling. The association between AP-1 transcriptional activity and clone size in aged donors connects to emerging literature implicating inflammatory programs in HSC aging^26,27^, and suggests that inflammatory regulatory states may confer a selective advantage contributing to clonal dominance.

In multiple myeloma, mtDNA phylogenies resolved tumor subclonal structure beyond the resolution of CNA analysis, revealing varying degrees of clonal remodeling in the three patients that corresponded to treatment response. Phylogeny–state analysis distinguished heritable from plastic tumor programs and nominated cell states associated with therapy resistance. These findings suggest that integrating mtDNA phylogenies with multiome profiling in larger cohorts can help identify clinically relevant drug resistance programs.

Our study reveals both the strengths and limitations regarding the temporal resolution of mtDNA-based lineage tracing. Over long timescales, mitochondrial variants undergo drift and can be lost, eroding phylogenetic signals in early branches. For instance, native human hematopoiesis is a massively polyclonal system, and sampled cells often share ancestry from early divisions (e.g., decades ago) rather than recent cell divisions. Indeed, mtDNA-based recovery of HSC clades depended strongly on branch time (measured by accumulated nuclear mutations): early splits (branches occurring before 100 mutations of molecular time, corresponding to the first few years after birth) were poorly recovered (∼2%), in contrast to recovery rates of up to 44% for later and more recent splits (J≥0.5, **Figure S5C**). Among clades correctly recovered, only 8/83 (∼10%; J≥0.75) were early splits, indicating that most HSC clades detectable by mtDNA mutations reflect postnatal expansions rather than developmental branching (**Figure S5D**). MitoDrift’s confidence-based refinement approach addresses this limitation by explicitly collapsing early splits with insufficient signal, focusing analysis on well-supported clades that share a common variant drift history. Consequently, our *in vivo* WGS-based benchmark demonstrates high precision for confident branches in the inferred phylogeny, though with more limited recovery of early ancestral structure.

Our findings suggest that integration of multiple endogenous lineage markers can further enhance tracing resolution in native human tissues. The complementary temporal resolution of nuclear and mitochondrial mutations motivates approaches that integrate both, where nuclear CNAs or SNVs support reconstruction of ancestral clonal expansions and mtDNA variants resolve more recent subclonal dynamics. Consistent with this, in donor MM1, CNA subclones — which delineate major lineage branches — were resolved as multiple clades in the mtDNA-based phylogeny with unclear relatedness to each other (polytomy), likely because no mtDNA variant reached sufficient heteroplasmy to mark that clonal divergence. Conversely, mtDNA-based phylogeny resolved fine-scale lineage relationships that copy number analysis alone could not distinguish, reflecting recent clonal diversifications that emerged after CNA acquisition. Beyond genomic variants, methylation-based epimutations may provide a higher resolution for recent clonal expansions before somatic mutations accumulate on mtDNA and drift to detectable heteroplasmy levels^23,41^.

In summary, we show that modeling of mtDNA inheritance dynamics enables precise lineage inference from single-cell multiome data. The MitoDrift framework establishes a foundation for quantitative lineage–state analyses in primary human tissues at scale, linking clonal history to transcriptional and epigenetic programs in development, aging, and disease.

## Methods

### Algorithm Overview

MitoDrift infers single-cell lineage trees from mtDNA variant allele counts using a discrete Wright–Fisher hidden Markov tree model (WF-HMT). For each mtDNA locus ℓ, the unobserved heteroplasmy state at every node *u* on the lineage tree is treated as a latent random variable, while observations are available only at the leaves (present-day cells) as finite mutant/reference molecule counts from single-cell sequencing. The model jointly specifies (i) a Wright–Fisher drift transition model along edges and (ii) a binomial observation model at leaves, enabling likelihood-based inference of both tree topology and model parameters.

#### State space

For each variant (locus) ℓ and each node *u* on the tree, the latent heteroplasmy state *z*_ℓ,*u*_ takes values on a discretized VAF grid {*p*_1_, …, *p*_*C*_} with *C* = *k* + 2 states: *k* internal bins over (0, 1) plus boundary states at 0 and 1 (default k = 20).

#### Wright–Fisher drift transitions

Along each parent to child edge, we model heteroplasmy drift as a Markov chain whose transition matrix *A* is a real-valued *C* × *C* matrix derived from a Wright–Fisher process with effective mitochondrial population size *N* over *g* generations. We first define a one-generation allele-count transition on counts *x* ∈ {0, …, *N*} as *T*_*x*→*y*_ = *Pr*(*Y* = *y*∣*X* = *x*) = *Binom*(*N*, *y*; *x*/*N*), where *N* is the population size. The *g*-generation transition is obtained by matrix power *T*^*g*^. We then aggregate allele-count probabilities into the discretized VAF bins to obtain *A* over {*p*_1_, …, *p*_*C*_}. In addition to drift, we model mutagenesis as rare events that introduce variants at homoplasmic boundaries. Let µ denote a boundary mutation rate that governs transitions from the homoplasmic states (VAF 0 or 1) into intermediate heteroplasmy states (and symmetrically in the reverse direction). Concretely, we modify the boundary rows of *A*: from the 0-VAF boundary (and symmetrically from 1-VAF), we inject a total probability mass µ into non-boundary states, with the remaining probability staying at the boundary. The root is fixed to the 0-VAF state (mutation absent at the root).

#### Observation model

For each locus ℓ and leaf cell *c*, we model the observed alternate count as 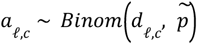 with 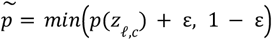, where *d*_ℓ,*c*_ is the total depth (total molecules) and ε is an additive error term that captures residual biases such as false positives/contamination and mapping artifacts. This emission model formalizes that observed VAFs are noisy, finite-sampling measurements of underlying heteroplasmy states.

#### Likelihood computation on a fixed tree

Given a topology, *A*, and leaf emission likelihoods, we compute the marginal likelihood for each locus by sum–product message passing (belief propagation) on the tree^14,15^. Because the graph is a tree, this calculation is exact. The total log-likelihood is the sum of per-locus log partition functions (normalizing constants).

Concretely, letting *T* = (*V*, *E*) denote a rooted tree over nodes *V* and edges *E*, and letting *D*_ℓ_ = {(*a*_ℓ,*c*_, *d*_ℓ,*c*_)}_*c*∈*leaves*_ denote the observed allele counts at locus ℓ, the likelihood factorizes across loci:

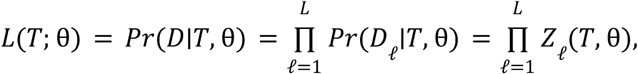

and thus 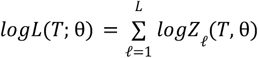 Each per-locus partition function marginalizes the latent heteroplasmy states *z*_ℓ,*u*_ ∈ {*p*_1_, …, *p*_*C*_} over all tree nodes:

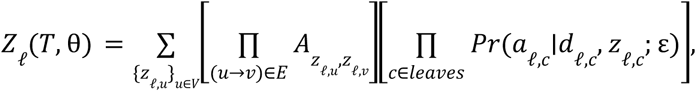

Where *Pr*(*a*_ℓ,*c*_ ∣*d*_ℓ,*c*_, *z*_ℓ,*c*_; ε) = *Binom a*_ℓ,*c*_; *d*_ℓ,*c*_, *min*(*p*(*z*_ℓ,*c*_) + ε, 1 − ε) under the observation model, and the root is fixed to the 0-VAF boundary state by a delta prior (mutation absent at the root).

#### Parameter fitting by expectation–maximization

Given an initial tree, we fit *g*, µ, and ε by expectation–maximization (EM), maximizing the observed-data likelihood under the WF-HMT.

- **E-step.** For each locus ℓ, we run belief propagation to compute posterior node marginals γ_ℓ,*u*_ (*c*) = *Pr*(*z*_ℓ,*u*_ = *p_c_* ∣*data*) and posterior edge marginals ξ_ℓ,(*u*→*v*)_ (*r*, *c*) = *Pr*(*z*_ℓ,*u*_ = *p*_ℓ,*u*_, *z* = *p*_*c*_ ∣*data*). We then accumulate expected transition counts across loci and edges,

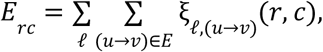 where *E* denotes the set of directed edges in the tree.
- **M-step.** We maximize the expected complete-data log-likelihood by separating the transition and emission terms. We update (*g*, µ) by bounded optimization of the transition objective

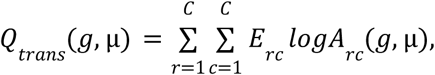 where *A*(*g*, µ) is the Wright-Fisher transition matrix with boundary mutation rate µ. For non-integer *g*, we use linear interpolation between adjacent integer-generation matrices,

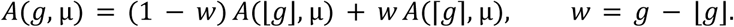

We update the emission error ε by 1D bounded optimization of

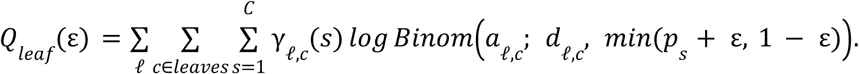

In practice, we optimize (*g*, *log*µ) using L-BFGS-B on − *Q*_*trans*_ with box constraints, and optimize *log*ε with a 1D bounded search on − *Q*_*leaf*_.

#### Tree topology inference and uncertainty estimation

We obtain an initial tree topology using neighbor joining (NJ) on pairwise Manhattan distances between cells computed from their observed VAF profiles. To quantify topological uncertainty and sample tree topologies consistent with the observed VAF data and drift history, we run Metropolis–Hastings Markov chain Monte Carlo (MCMC) over tree topologies with random NNI proposals. Proposals are accepted or rejected based on the log-likelihood difference under the WF-HMT model, yielding posterior clade support (the frequency of each bipartition across sampled trees).

Naively recomputing the full WF-HMT likelihood after each NNI proposal requires re-running belief propagation across the entire tree for every locus, which becomes prohibitively expensive for thousands of cells. We therefore exploit that NNI is local and reuse cached dynamic-programming quantities when evaluating proposed topologies. Our belief-propagation implementation aggregates information upward from leaves to the root. For each locus ℓ and node *v*, define a length-*C* vector *F*_ℓ_ [*v*, ·] as the child-to-parent contribution that is added to the parent’s accumulator (i.e., the quantity contributed by *v* to its parent after summing over *v*’s latent state). For a parent node *p* with children *a* and *b*, the “children message” accumulator at *p* is the elementwise sum of child contributions,

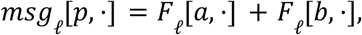

and the root partition function for locus ℓ is

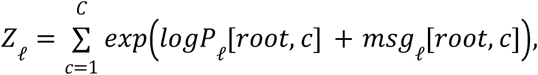

where *logP*_ℓ_ [*root*, *c*] is the leaf/internal-node likelihood term (from the binomial emission at tips; 1 at internal nodes) expressed on the discretized VAF grid.

To compute *F*_ℓ_ [*v*, ·] efficiently across loci, we form, for each state *c*, a stabilized vector

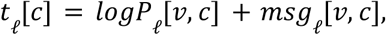

compute per-locus maxima *m*_ℓ_ = *max*_*c*_ *t*_ℓ_ [*c*], and set *u*_ℓ_ [*c*] = *exp*(*t*_ℓ_ [*c*] − *m*_ℓ_). Then, for each parent-state *r*, the child contribution can be written as

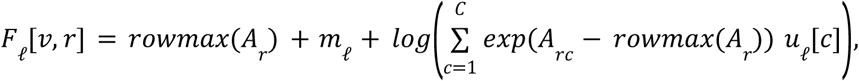

where *A* is the (log) Wright–Fisher transition matrix, and *rowmax*(*A*_*r*_) is the maximum element in row *r* used for numerical stabilization.

During topology MCMC, we cache *F*_ℓ_ [*v*, ·] for all nodes *v* and loci ℓ for the current tree, as well as the current total log-likelihood 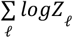 Consider an NNI proposal at an internal edge (*p*_1_ → *p*_2_); this swap changes only the assignment of one child subtree between *p*_1_ and *p*_2_. All cached *F* values strictly within the unaffected subtrees remain valid. We recompute *F* for *p*_2_ and *p*_1_ under the proposed child configuration, then walk upward along the unique path from *p*_1_ to the root, recomputing *F* only for nodes on that path (each time combining the updated child contribution from below with the unchanged sibling subtree). Once the root is reached, we recompute the proposed 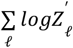 using the updated root children contributions and obtain the Metropolis–Hastings acceptance decision from the log-likelihood difference.

This yields an update cost proportional to the height of the affected path, *O*(*height* × *L* × *C*^2^) (with efficient vectorization across loci), rather than *O*(*n* × *L* × *C*^2^) recomputation over all *n* cells.

#### MCMC mixing diagnostics

We monitored mixing across parallel topology MCMC chains using an average standard deviation of split frequencies (ASDSF) computed against a fixed set of clades defined by the initial neighbor-joining tree topology. After each batch of MCMC iterations, we computed clade frequencies in each chain using all samples accumulated so far. For each target clade *k* and chain *c*, we define *f*_*c*,*k*_ as the fraction of sampled trees in chain *c* that contain clade *k* (rooted clade matching). We then compute the across-chain standard deviation for each clade, *SD*_*k*_ = *sd*_*c*_ (*f*_*c*,*k*_), and define ASDSF as the average, across the target clade set, of the between-chain variability in those clade frequencies. Intuitively, if different chains have converged to the same posterior over topologies, then they should assign similar frequencies to each target clade and the ASDSF will be small. We can optionally use a prespecified ASDSF threshold as an automatic stopping criterion, terminating topology sampling once the batch-wise ASDSF falls below this threshold.

### Confidence-based topology refinement and cutoff selection

To increase precision in the inferred lineage tree and prioritize well-supported clades, we refined the initial NJ tree topology by collapsing branches with posterior support below a chosen cutoff (τ). A stringent threshold (high τ) can discard biologically meaningful clonal structure, whereas a lenient threshold (low τ) can retain spurious splits not supported by the underlying variant data. Because different samples vary in mtDNA mutation density, cell number, and technical noise, we selected dataset-specific confidence thresholds using a parameter sweep over candidate values of τ. At each cutoff, we evaluated concordance between retained tree clades and individual variant carrier sets (cells exceeding a VAF threshold for a given variant) by Jaccard similarity, yielding a variant-based precision–recall diagnostic without external ground truth: precision measures the fraction of retained clades that match a variant carrier set, while recall measures the fraction of variants recovered by a retained clade above the cutoff. This diagnostic guides threshold selection by identifying τ that balance clade retention against poorly-supported splits. Diagnostic curves and chosen cutoffs for representative datasets are reported in **Figure S2** and **Table S1**. Because individual variant carrier sets are affected by dropout, contamination, and homoplasy, they are imperfect proxies for true clonal structure; the variant-based precision–recall curve therefore provides a relative diagnostic for comparing cutoffs within a dataset rather than an absolute measure of tree accuracy.

For the regulon heritability analysis in Young1 HSCs, where the sample is highly polyclonal and most true clones are small, we used an alternative size-aware topology refinement criterion that preferentially retains fine-scale structure near the tips of the tree. Fixed-*τ* thresholding controls per-clade error probability uniformly across the tree; in polyclonal samples, this can collapse the fine-scale tip-proximal splits that carry the lineage signal relevant for heritability estimation. The alternative criterion instead controls the expected number of mis-assigned cells: for each internal node with clade size *n*_*c*_ and posterior support *p*, the expected number of mis-assigned cells is (1 − *p*) *n*_*c*_. We collapsed branches where this quantity exceeded ɛ · *n*_*tip*_, where *n*_*tip*_ is the total number of tips and ɛ is a user-specified tolerance. Because (1 − *p*) *n*_*c*_ is naturally small for small clades near the tips, this criterion retains fine-scale structure while collapsing deeper, larger groupings whose uncertainty would propagate to many cells.

### Defining clones using the refined lineage tree

In downstream analyses that require a discrete clone partition, we define “clones” operationally from the refined tree as the set of top-level subtrees descending from the root: each child of the root defines one top-level clade consisting of all its descendant tips. Each such clade is the maximal group of cells whose internal branching relationships are all supported above the confidence threshold—any larger grouping would require traversing an edge that was collapsed. We treat each multi-cell top-level clade as one detected clone at the chosen cutoff. Tips that attach directly to the root after collapsing low-confidence branches (“root singletons”) should not be interpreted as biologically independent clones; rather, they reflect cells whose lineage relationship to other sampled cells is not confidently resolved at cutoff τ. This definition reflects the structure of the mtDNA lineage signal. Within each top-level clade, cells share enough distinctive mtDNA heteroplasmy variation—accumulated through intracellular drift in a shared ancestor—to support confident grouping above the chosen threshold.

Between top-level clades, the deeper branching order is not confidently resolved: the root polytomy is a soft polytomy reflecting limited phylogenetic signal, not a claim of simultaneous divergence. The number of resulting clones and their boundaries are therefore functions of both the underlying clonal structure and the confidence threshold τ: stricter cutoffs yield fewer, more conservative clones (plus more unresolved cells), while lenient cutoffs retain finer-grained structure at the cost of reduced per-clade precision.

### Multiple myeloma patient bone marrow samples

Bone marrow samples from multiple myeloma patients were ordered from Discovery Life Sciences. According to the provider, samples were collected under an IRB-approved sample-banking protocol with informed consent.

| Patient ID | Age | Gender | Treatment |
| --- | --- | --- | --- |
| MM1 | 74 | Male | Cy/Bor/Dex 6 cycles+ Radiotherapy |
| MM2 | 82 | Male | Cy/Bor/Dex 6 cycles |
| MM3 | 70 | Female | Cy/Bor/Dex 8 cycles |

### ReDeeM workflow

ReDeeM was performed as previously described. Briefly, ReDeeM is a modified 10x Genomics droplet-based single-cell multiome protocol in which intact cells are fixed and mildly permeabilized to maximize mitochondrial DNA (mtDNA) retention. mtDNA fragments are then selectively enriched by hybrid-capture using mtDNA-specific probe sets and sequenced at high depth with unique molecular identifiers (UMIs) to enable error-corrected mtDNA mutation calling. In parallel, matched scRNA-seq and scATAC-seq libraries are prepared from the same droplets using the 10x workflow, with all three libraries linked by shared single-cell barcodes for joint phylogenetic and cell-state analyses.

### Lentiviral barcoding with ReDeeM

To benchmark MitoDrift’s mtDNA-based lineage reconstruction against exogenous clonal ground-truth, we used a lentiviral static barcoding system (LARRY v2)^16^ with primary human HSCs. Briefly, healthy donor HSPCs (CD34+) were transduced with the LARRY lentiviral barcode library, and 1,000 LARRY-positive HSCs/MPPs (CD34^+^, CD45RA^-^, CD90^+^) were sorted and plated for *in vitro* expansion and differentiation. Following 10 days of ex vivo culture, 150,000 expanded progeny cells were profiled using the ReDeeM protocol as described above with an additional targeted capture of LARRY barcodes from the matched cDNA library, which determines ground-truth clone membership and enables direct comparison between mtDNA-inferred lineages and LARRY barcode-defined clones.

### Lentiviral barcode data preprocessing

We processed raw FASTQ reads with the bartab pipeline (Nextflow, DaneVass/bartab v1.4) in single-cell mode using known flanking constants (up: CGATTCCAGT; down: ATCGCCACCG) and a fixed cell barcode/UMI pattern (16C+12N). A curated whitelist derived from bulk-DNA level barcode was applied at parse time to restrict extraction to valid barcodes. Reads were mapped to the provided reference, barcodes and UMIs were parsed and deduplicated, and per-cell barcode count tables were written to the output directory. Static LARRY barcode clone assignments were derived from the barcode count table using a graph-based quality control procedure. First, barcode-cell associations were filtered to retain only entries with UMI count >= 3, and cells carrying more than 3 distinct barcodes were excluded as likely multiplets. Cells were then connected in an undirected graph where an edge between two cells indicates they share at least one barcode. To remove cells that spuriously bridge otherwise distinct clones, nodes with betweenness centrality above the 95th percentile were removed. After recomputing connected components, cells were further filtered by requiring an intra-component connectivity ratio (node degree divided by component size) of at least 0.5. This step removes loosely connected cells at the periphery of components that may represent barcode cross-contamination or doublets. The remaining connected components define the final clone assignments, with each component representing a clonal group sharing one or more static barcodes. Singleton clones (component size = 1) were excluded from downstream analyses.

### Single-cell multiome preprocessing and embedding

We used *Seurat* (v5.3.0) and *Signac* (v1.14.0) for multiome preprocessing. Cells with fewer than 300 detected genes were removed prior to doublet detection. Doublets were identified using *scDblFinder* (v1.22.0) on the RNA count matrix with cluster-aware simulation and removed. Quality control filtering then removed cells with >15,000 RNA features, >20% mitochondrial reads, <2,000 or >30,000 ATAC features, TSS enrichment <1.5, or nucleosome signal >2. Cell type labels were assigned by projecting query cells onto a BoneMarrowMap Symphony reference. Hematopoietic cell types were predicted by k-nearest neighbor classification against the reference.

For joint RNA/ATAC analyses (e.g., wnnUMAP), we used a weighted-nearest-neighbor (WNN) workflow that (i) performs RNA normalization with *SCTransform* (v0.4.2), computes RNA PCA, (ii) performs ATAC TF-IDF normalization, selects top features with FindTopFeatures(min.cutoff = "q0"), computes ATAC LSI with RunSVD, (iii) constructs a joint neighbor graph with FindMultiModalNeighbors (RNA dims 1:30; ATAC dims 2:30), and (iv) clusters cells on the WNN graph (FindClusters(graph.name = "wsnn", algorithm = 1)) and computes a 2D embedding with RunUMAP(nn.name = "weighted.nn").

### ReDeeM-based variant calling

For primary mtDNA variant detection, we used the redeemV (https://github.com/chenweng1991/redeemV) and redeemR (https://github.com/chenweng1991/redeemR; v2.0) pipeline^4^ and filtered the resulting variant calls using the Filter-2 procedure with edge trimming set to 9 base pairs as previously described^42^. Briefly, ReDeeM Filter-2 uses the same consensus and downstream filtering as the original redeemV pipeline, with two updates in redeemR. (1) It adds a fragment-end proximity filter: after consensus error filtering, each mutation is annotated by its distance to the nearest fragment end and variants within d bp are removed (d = 9 is used). (2) It replaces the original hard inclusion rule (max allele ≥ 2 in any cell and detected in ≥ 2 cells) with a binomial noise model: residual post-consensus noise is assumed binomial, and variants are filtered if their across-cell pattern is consistent with the binomial null (χ² test; adjusted *P* > 0.05). The Filter-2 pipeline can be accessed through the redeemR package.

### Benchmarking the impact of variant calling on lineage reconstruction performance

To evaluate the impact of different upstream mtDNA variant calling strategies on phylogenetic reconstruction, we evaluated the precision-recall against the lentiviral barcoding (LARRY) ground truth using the MitoDrift framework. We compared two ReDeeM mutation filtering regimes as previously described Filter-1^4^, Filter-2^42^, and more stringent filtering approaches with additional heteroplasmic variant allele frequency cutoffs on Filter-2, removal of all variants supported by one molecule, as well as a previously published mtDNA mutation calling approach (mgatk; v0.7.0). Precision-recall benchmarking identified ReDeeM Filter-2 as the best-performing strategy for downstream phylogenetic reconstruction, with Filter-1 also showing high performance (**Figures S13,14**). Both approaches outperformed more stringent filtering regimes, including mgatk-like settings and increasingly strict heteroplasmy cutoffs. Notably, removing all 1UMI-supported variants reduced performance, indicating that this class contains genuine lineage information, providing more signal than noise. We also observed a precision ceiling under stringent filtering: with aggressive filtering, precision did not improve proportionally, while recall declined, consistent with depletion of informative low-VAF variants that are required for clone resolution (**Figure S13**). Consequently, we selected ReDeeM Filter-2 as the default filtering strategy for the remainder of the study.

For the benchmarking, we compared lineage reconstruction precision-recall performance across predefined filtering strategies in ReDeeM and mgatk using a matched evaluation framework. For ReDeeM, we analyzed S consensus with Filter-1 (keep consensus-called mutations that recur across cells and reach ≥2 eUMIs in at least one cell), Filter-2 (Filter-1 + fragment edge-trimming + binomial/FDR test to remove remaining noise), Filter-2 with no-1UMI (mutation observations with 1-UMI support are removed), and Filter-2 with additional heteroplasmy thresholds (≥ 0.02, 0.04, 0.06, 0.08, 0.10); for mgatk, we used fr2_mc3 preprocessing (minimum 2 supporting reads per strand; variant must be detected in at least 3 confident cells; https://github.com/caleblareau/mgatk) with baseline or additional heteroplasmy thresholds ≥ 0.07 settings^43^. Ground-truth cell-clone annotations were derived from curated barcode metadata (clone size ≥2). We generated 10 fixed clone-based subsets and reused the same subset definitions for every condition to control for composition effects, then inferred one MitoDrift tree per condition-subset with matched inference parameters.

For evaluation, as described, recall was defined as clone-level recovery against ground truth across a MitoDrift branch support *τ* sweep, and precision was computed from inferred clade-to-clone matching at a fixed clone-truth overlap threshold (Jaccard 0.5 or 0.75). In addition to end-to-end precision-recall (same set of cells for both precision and recall across all conditions), we also used conditional precision, defined as precision after restricting evaluation to evaluatable cells (excluding cells without mutation after filtering in the precision branch), while retaining full-cell recall. Per-subset curves were aggregated by averaging across the 10 matched subsets at each *τ*.

### MAESTER variant calling

For the MAESTER dataset of a donor with clonal hematopoiesis^3^, we followed the variant calling, filtering, and quality control procedures described in the original publication. For final variant selection, we required at least 5 cells with VAF exceeding 5%.

### Single-colony WGS variant calling

For the Mitchell et al. single-colony WGS dataset, we used GATK Mutect2 (v4.6.2.0) to call variants (genome version: GRCh37/hs37d5). To identify a conservative set of mtDNA variants for downstream phylogeny inference, we defined a variant as *confidently detected* in a colony using a one-sided binomial test on alternate counts (*a*) out of total depth (*d*), with a background error rate of *p* = 0. 01 (alternative = greater). For each variant, we applied Bonferroni correction across colonies and called confident detection when adjusted *P* < 0.05. We retained variants with confident detection in at least two colonies. We then removed likely germline or fixed variants using the global VAF across all colonies (∑*a*/∑*d* > 0. 75), and filtered recurrent low-VAF/high-prevalence sites by excluding variants with low mean VAF among detected colonies (mean VAF < 0.05 among colonies with VAF > 0.002) combined with broad prevalence (present in >25% of colonies). Because donor KX003 was found to have a germline polymorphism^9^, we applied a correction for variant 16519 T>C by re-assigning reference and alternate alleles based on the donor’s germline genotype.

### mtDNA-based clonal diversity

We compute mtDNA-based Shannon Diversity Index (mtSDI) by applying the Shannon diversity index to the resulting clone partition: letting *n* denote the number of cells in clone *k* and 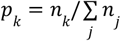, we compute 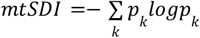, counting each root singleton as a size-1 unit. Under this convention, mtSDI should be interpreted as an upper bound on detectable clonal diversity at the chosen cutoff, because unresolved cells are maximally separated rather than merged; mtSDI therefore depends on mtDNA marker density and sequencing sensitivity in addition to the underlying clone-size distribution. Because mtSDI depends on branch confidence estimates, which are influenced by MCMC parameters and cell count, we controlled for these factors to ensure comparability within each analysis. For donor-level mtSDI in the WGS benchmark cohort, we inferred a single MitoDrift tree per donor over all cells using identical MCMC parameters and confidence threshold (τ = 0.05) across donors and computed mtSDI from the resulting clone partition. For cell-type-specific comparisons across donors, we randomly subsampled 200 cells per cell type per donor and inferred a MitoDrift tree independently for each subsample using identical MCMC parameters and branch confidence threshold (τ = 0.01). Cell types with fewer than 200 cells in a given donor were excluded. We repeated this procedure across 20 random subsamples per donor–cell type pair and computed mtSDI from the resulting clone partition for each replicate, reporting the mean and 95% confidence intervals across replicates. For the donor-multiplexed ReDeeM cohort, which yields fewer cells per donor due to sample multiplexing, we subsampled 45 cells per cell type per donor across 10 random replicates, and applied confidence refinement (τ = 0.01). Cell types were grouped coarsely to ensure sufficient cell counts for each lineage category. For the MAESTER clonal hematopoiesis dataset and the multiple myeloma dataset, we inferred a single MitoDrift tree per sample over all cells (across cell types or treatment timepoints, respectively), applied confidence refinement (τ = 0.125 for MAESTER, τ = 0.075 for MM samples), and then subset the refined tree by cell type or timepoint to compute per-partition mtSDI. This approach ensures comparability within a sample because all partitions derive from the same underlying tree and model parameters.

### Tree accuracy metrics and clade matching

To quantify concordance between inferred and ground-truth trees (or clone partitions), we represented each internal node as a clade (the set of descendant cells/tips) and compared clades across two trees based on tip overlap. We summarized agreement using Jaccard similarity (intersection over union) between clades and evaluated trees at a specified matching threshold (e.g., Jaccard ≥ 0.5).

Because inferred lineage trees can be polytomous (for example after collapsing weakly supported edges), evaluation must reflect the tradeoff between the number of predicted branches and the accuracy of those branches. Conceptually, we define: (i) recall as the number of correctly predicted (matched) ground-truth clades divided by the total number of ground-truth clades, and (ii) precision as the number of correctly predicted (matched) inferred clades divided by the total number of inferred clades, using the same clade-matching criterion.

When the ground truth is defined by static lentiviral barcodes (LARRY), we convert the barcode clone assignment into a polytomous “barcode tree” in which each barcode corresponds to one branch/clade: a single root connects directly to one internal node per barcode clone, and all cells carrying that barcode are attached under that node. This representation enables clade-level matching between inferred phylogenies and barcode-defined true clones using the same overlap-based criteria.

### Comparison with existing methods

We compared MitoDrift to several baseline lineage reconstruction approaches in our benchmarking analyses. We included distance-based phylogenetic methods (NJ and UPGMA), a perfect-phylogeny (infinite-sites) approach (MERLIN; https://github.com/raphael-group/MERLIN), a non-phylogenetic approach (Signac; v1.14.0), and branch confidence estimation by Transfer Bootstrap. For the bootstrap baseline, we generated bootstrap replicate trees by resampling the variant matrix and computed bootstrap support values onto the NJ topology using *phangorn::transferBootstrap*; we then evaluated trees after collapsing clades below a range of support thresholds to yield polytomies comparable to confidence-refined MitoDrift trees. For the Signac baseline, we used Signac’s *FindClonotypes* procedure to directly cluster cells into discrete clonotypes based on the VAF matrix. Briefly, we computed a k-nearest-neighbor graph (k = 10; cosine distance) and corresponding SNN graph, and performed graph-based community detection (SLM algorithm) at a range of resolution values. Because Signac returns a clonotype assignment rather than a tree topology, we converted each clonotype assignment into a polytomous "clone tree" (one internal node per clonotype under the root).

For the lentiviral barcoding benchmark, we randomly subsampled 45 clones (defined by LARRY barcodes) per replicate and repeated this procedure for 10 replicates. For each replicate, we inferred trees with each method and computed clade precision and clone recall using the clade-matching framework described above. Precision–recall curves and summary statistics were obtained by averaging precision and recall across the 10 technical replicates at each method-specific cutoff value. For the single-colony WGS benchmark cohort, we randomly subsampled 150 cells per sample per replicate and repeated for 10 replicates. For each replicate, we evaluated mtDNA-based trees against the corresponding nuclear SNV phylogeny using the same clade-level precision/recall framework, and averaged precision and recall across replicates.

To benchmark clonal diversity estimation using the pLARRY dataset, we generated 200 simulated cell sets spanning a wide range of barcode-defined SDIs by subsampling cells at the clone level (barcode-defined clones). For each replicate, we formed a subset of approximately 300 cells by selecting whole barcode clones (not individual cells) via size-biased sampling without replacement (weights 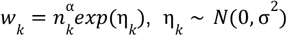) and adding clones until reaching 95% of the target size; sweeping α evenly from + 3 to − 3 varied the clone-size skew from oligoclonal to polyclonal. We computed ground-truth SDI from barcode assignments and mtSDI from the corresponding confidence-refined MitoDrift clone partition and compared SDI estimates across replicates.

### Lineage–state coupling analysis with PATH

To quantify coupling between lineage and cell state, we used PATH (v1.0) to conduct phylogenetic association test on a rooted lineage tree^19^. We computed a tip-by-tip weighting matrix *W* derived from the tree and used it to compute a weighted correlation between feature values across related cells, reporting a Z-score for each pairwise state comparison (or feature) and computing two-sided P values from a normal approximation followed by Benjamini–Hochberg correction. In our analysis, *W* was constructed using a Brownian-motion covariance (*ape::vcv* after setting all branch lengths to 1 and zeroing the diagonal with row normalization).

### Progenitor state-to-fate regression

To link early progenitor regulatory state to downstream lineage output in the mtscATAC-seq CD34+ differentiation dataset^3^, we focused on day 8 progenitor cells and defined a per-cell fate-bias score using the inferred mtDNA phylogeny. We subset the confidence-refined MitoDrift tree to day 8 progenitors (including progenitor subtypes) and day 14/20 differentiated cells. For each day 8 progenitor cell, we computed the composition of the *k* = 5 nearest non-progenitor neighbors by patristic distance (assuming unit branch lengths), restricting the neighbor pool to differentiated (myeloid or erythroid) cells. We defined *myeloid bias* as the fraction of these neighbors annotated as myeloid.

To identify transcription factor programs associated with fate bias, we performed per-feature linear regression of chromVAR TF activity scores in day 8 progenitors on the myeloid-bias score while controlling for sequencing covariates (total depth and mtDNA coverage): *y* ∼ *x* + *depth* + *mtDNAcoverage*, where *y* is the chromVAR score for a given TF and *x* is the myeloid-bias score. We then ranked the TFs by their regression coefficient for *x*.

### Clade enrichment scores

To visualize enrichment of traits (e.g., treatment timepoints, cell states, or continuous scores such as pseudotime) across the phylogenetic tree, we computed z-scores quantifying deviation of each clade from the global trait distribution. For each internal node, we computed the mean trait value among its descendant tips and compared this to the global mean across all tips. The z-score was calculated as the deviation from the global mean divided by a size-adjusted standard error, where the standard error equals the global standard deviation divided by the square root of the clade size. This formulation accounts for the expected reduction in variance for larger clades while detecting clades that are significantly enriched or depleted for the trait of interest. Two-sided *P* values were computed from the normal distribution and corrected for multiple testing using the Benjamini–Hochberg procedure, restricting tests to clades above a minimum size threshold. For binary traits such as treatment timepoint, tip scores were encoded as indicator variables (0 or 1); for continuous traits such as pseudotime or regulon activity, tip scores were used directly. To facilitate tree visualization, we propagated z-scores from internal nodes to their descendants by assigning each branch the maximum absolute z-score encountered along the path from the root to that node, allowing branches to be colored by the enrichment of their most significant ancestral clade.

### Regulon heritability

For regulon analyses in young and aged donor HSPCs profiled by ReDeeM, we used SCENIC+ (v1.0a2) to perform gene regulatory network inference and compute regulon activity scores (AUCell) for RNA and ATAC modalities. To reduce confounding from sequencing depth, we generated depth-adjusted regulon scores by regressing out total RNA counts (for RNA-based scores) or total ATAC counts (for ATAC-based scores) using Seurat and using the residualized scores for downstream phylogenetic tests. For regulon heritability analysis in the LARRY dataset, we used pySCENIC^44^ (v0.12.1) to infer gene regulatory networks and compute regulon activity scores from the RNA modality, as the ATAC library was not available for this dataset. We tested for phylogenetic signal (heritability) at each timepoint using the Cmean statistic with permutation-based *P* values using the *phylosignal* R package (v1.3.1). To identify regulons with statistically significant heritability across the two sampling timepoints in Young1 HSCs, we refined the tree topology using the expected number of misassigned cell criteria (ɛ < 0.002) and combined per-timepoint *P* values from the Cmeans test for each regulon using Fisher’s method, then applied Benjamini–Hochberg correction across regulons to obtain Q values.

### AP-1 regulon regression versus clone size

To quantify associations between inferred clonal expansion and regulatory programs, we regressed AP-1–associated regulon activity against inferred clone size within HSCs and MPPs. We used confidence cutoffs τ = 0.005 for Aged 1 and τ = 0.01 for Aged 2. Clone size was defined from the refined lineage tree as the number of descendant cells assigned to each clone.

For each regulon/feature, we fit a linear model of the form *y* ∼ *x* + *batch* + *covariates*, where *y* is the per-cell regulon activity, *x* is the per-cell clone size, and batch is an indicator variable capturing sample/batch membership (in this case, the HSC/MPP cell type label). Additional covariates (including ATAC and RNA library depth) were included as additive terms. We controlled for multiple testing across features using Benjamini–Hochberg to obtain adjusted *P* values (*Q* values).

### Copy-number inference in multiple myeloma dataset

To identify malignant plasma cells and quantify subclonal CNA structure in the myeloma multiome cohort, we performed CNA inference using Numbat (v1.5.2) in multiome mode, which leverages both RNA and ATAC coverage signal within the same cells^30,45^. Briefly, we generated per-cell allele counts for each donor and timepoint using Numbat’s pileup and phasing workflow, and constructed binned coverage matrices for each modality by aggregating RNA counts and ATAC fragments into fixed genomic bins (220 kb bins). We then ran joint multiome Numbat inference per donor across the two timepoints (pre-treatment and post-treatment), using Numbat parameters k=5, gamma=5, min_LLR=30, min_genes=5, t = 1e-5, max_nni = 100, and multi_allelic=FALSE. We built the reference expression profile by aggregating merged multiome binned counts (RNA+ATAC) across non-plasma broad cell types from all samples.

We used Numbat’s per-cell CNA posterior probabilities and clone assignments for downstream visualization and annotation of tumor burden and subclonal structure. To classify cells as tumor or normal, we used Numbat’s posterior compartment assignments. In donor MM1 where we performed detailed lineage-state analysis, we applied a k-nearest-neighbor smoothing procedure in the joint RNA/ATAC embedding space to reduce noise in single-cell clone assignments. For each cell, we identified its 50 nearest neighbors in the WNN UMAP embedding and reassigned the cell to the majority CNA clone among its neighbors, yielding refined clone assignments.

### Malignant plasma cell-state definition

To define cell states among malignant plasma cells in the myeloma dataset, we subset Seurat objects to tumor cells (identified via Numbat CNA inference) and performed graph-based clustering on the precomputed weighted nearest-neighbor WNN graph using the Louvain algorithm (resolution = 0.7). Clusters were labeled by timepoint to track state composition across treatment. For each cluster, we identified marker genes and transcription factor motifs using Wilcoxon rank-sum tests on RNA expression and chromVAR TF deviation scores, respectively. For subclonal cell-state analysis within CNA-defined tumor subpopulations (e.g., the 1q-gain subclone), we subset cells to the subclone of interest and recomputed the WNN graph on the reduced cell set before reclustering (resolution = 0.5). This approach enabled identification of cell states specific to genetically defined tumor subpopulations that were obscured in the full tumor analysis. To visualize phylogenetic relatedness to post-treatment cells at continuous resolution, we reclustered the 1q-gain subclone at fine resolution (resolution = 3.0), computed mean pairwise phylogenetic distances between each fine-grained pre-treatment cluster and post-treatment cells, and projected these community-level distances onto the cell-state embedding.

### Inter-community phylogenetic distances

To quantify phylogenetic relatedness between groups of cells (e.g., communities or timepoint-specific cell states), we computed pairwise tip-to-tip phylogenetic distances from the inferred tree (setting all branch lengths to 1) and summarized between-group distances using *picante* (v1.8.2) on a group-by-tip incidence matrix. When confidence intervals were required, we performed bootstrap resampling within each group by multinomial resampling of the group’s tip counts and recomputed community distance across bootstrap replicates to obtain empirical confidence intervals.

### Analysis software

Tree-based analyses and metrics were implemented in R (v4.4.2) using standard phylogenetics and single-cell analysis packages.

### Data availability

This study used the following previously published datasets: the mtscATAC-seq CD34+ differentiation dataset^2^ (GEO accession GSE142745), the MAESTER dataset^3^ (GEO accession GSE182685), the single-colony WGS cohort^7^ (EGA accession EGAD00001007851), and the ReDeeM healthy hematopoiesis dataset^4^ (GEO accession GSE219015). Processed sequencing data of Lentiviral barcoding ground-truth datasets for benchmarking mtDNA-based lineage tracing is available on figshare (https://doi.org/10.6084/m9.figshare.31323481). The raw and processed sequencing data of longitudinal multiple myeloma multiome will be available upon publication.

### Code availability

The MitoDrift software package is available on GitHub (https://github.com/sankaranlab/mitodrift). The analysis scripts and notebooks reproducing the figures and analyses in this paper will be available upon publication.

## Acknowledgements

We thank members of the Sankaran and Weissman labs for valuable comments. We thank Adam Sperling for helpful discussion regarding multiple myeloma biology. This work was supported by the Howard Hughes Medical Institute (V.G.S. and J.S.W.), the Mathers Foundation (V.G.S. and J.S.W.), the Manton Cell Discovery Network at Boston Children’s Hospital (V.G.S.), the Alex’s Lemonade Stand Foundation (V.G.S.), and National Institutes of Health (NIH) grants R01DK103794, R01CA265726, R01CA292941, R33CA278393, and R01HL146500 (V.G.S.). T.G. is a Howard Hughes Medical Institute Fellow of the Damon Runyon Cancer Research Foundation (DRG-2552-25). C.W. is supported by NIH Pathway to Independence Award (K99HG013991). L.I.Z., J.S.W., and V.G.S. are investigators of the Howard Hughes Medical Institute.

## Author contributions

T.G., C.W., J.S.W., and V.G.S. conceived the project and directed the studies. T.G. conceived and developed the MitoDrift algorithm and performed computational analyses. C.W. directed the experimental studies, performed experiments and computational analyses, and coordinated data collection. J.G., I.J., A.D., L.B., Y.S., E.M., and L.I.Z. assisted with experiments and data collection. M.P., E.K., C.R., and R.T. assisted with data analysis. J.S.W. and V.G.S. supervised the study. All authors contributed to writing the manuscript.

---

**Figure S1.**
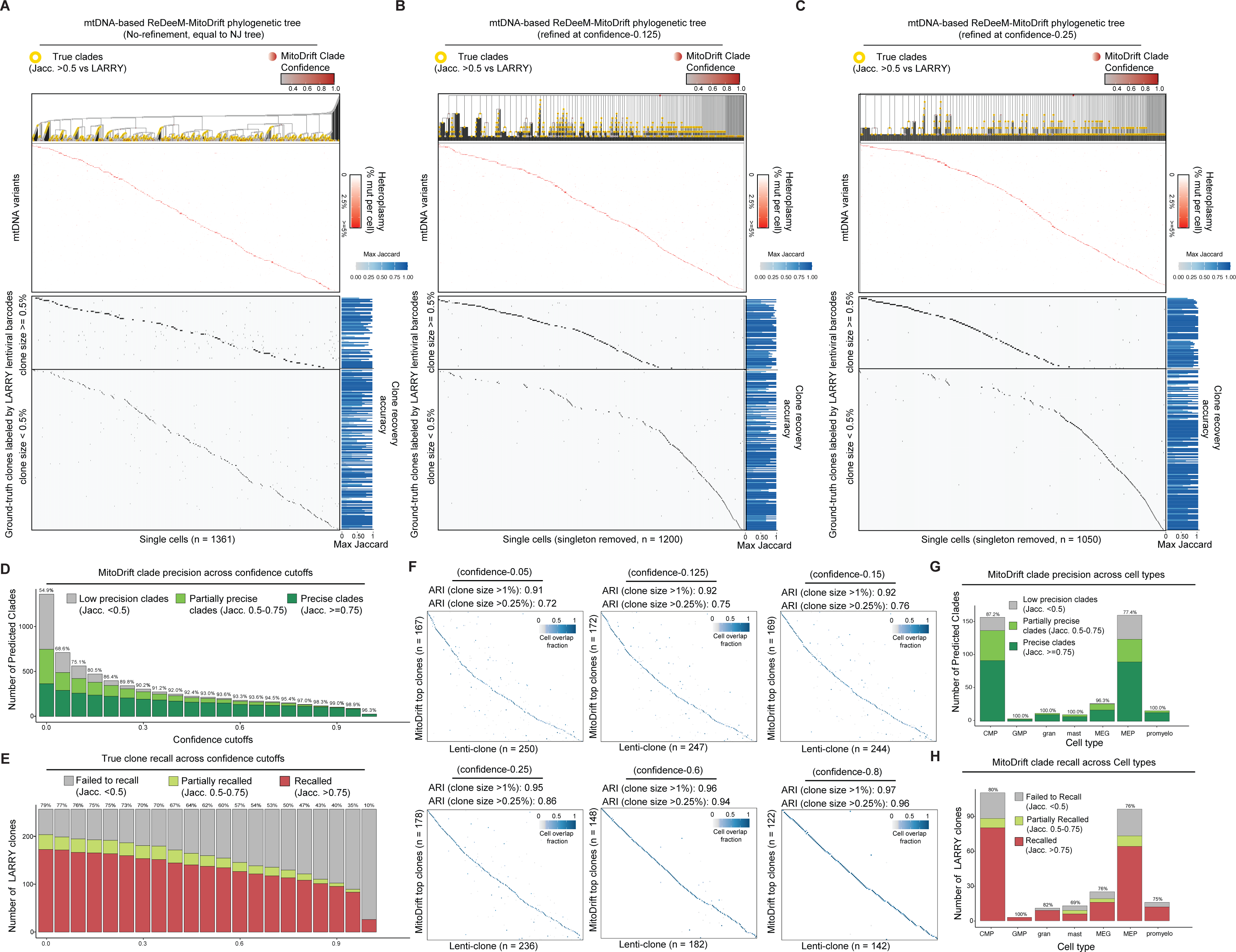
Clade precision and clone recall in the LARRY benchmark. (A–C) mtDNA-based ReDeeM-MitoDrift phylogenetic trees for the LARRY benchmark shown without confidence refinement (A) or refined at confidence thresholds τ = 0.125 (B) and τ = 0.25 (C). Trees are annotated with true clades based on overlap with LARRY barcode clones (Jaccard ≥ 0.5; yellow) and posterior clade confidence (color bar); heatmaps show mitochondrial variants across cells (top) and LARRY barcode-defined ground-truth clone identities (bottom), with per-barcode clonal recovery accuracy summarized as the maximum Jaccard overlap (right). (D) Precision of inferred clades with different confidence cutoffs, stratified by clade–clone overlap (false positives: Jaccard < 0.5; true positives: Jaccard 0.5–0.75 and ≥0.75). (E) Number of LARRY barcode clones recovered at each cutoff, stratified by best-matching inferred clade overlap (not recovered: Jaccard < 0.5; recovered: Jaccard ≥0.5 and ≥0.75). (F) Concordance between MitoDrift root-clone assignments and LARRY clone identities across confidence cutoffs, visualized as clone–clone overlap matrices (color indicates cell overlap fraction) and summarized by adjusted Rand index (ARI). (G) Clade precision stratified by annotated cell type. (H) Clone recall stratified by annotated cell type.

**Figure S2.**
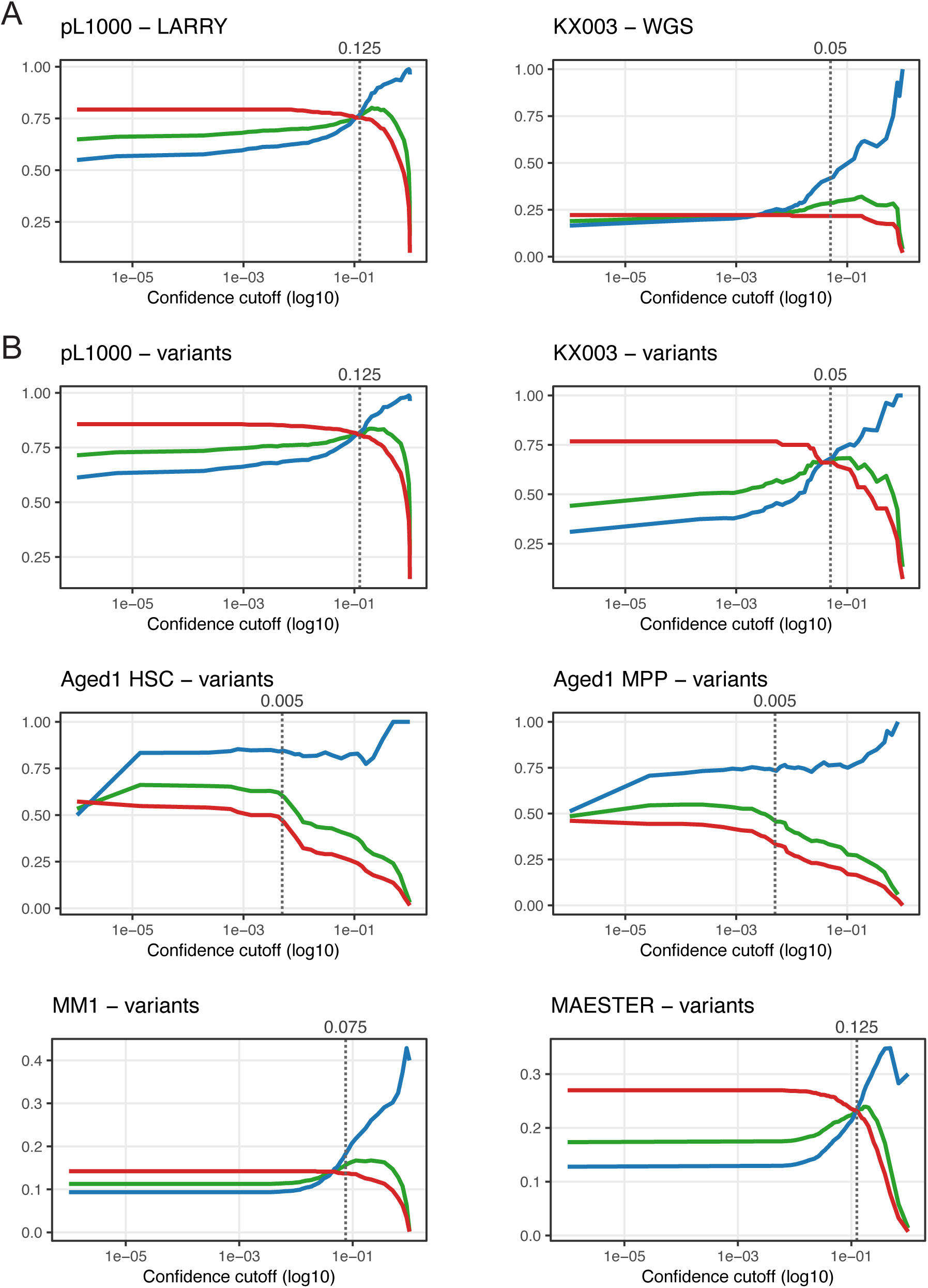
Confidence-cutoff diagnostic curves across datasets. (A) Precision, recall, and F1 score as a function of the posterior clade-support cutoff (τ; x-axis, log10 scale) when evaluating confidence-refined MitoDrift tree against external ground truth in LARRY (pL1000; barcode clones) and the WGS benchmark (KX003; nuclear SNV phylogeny clades). (B) For datasets without external ground truth, an internal diagnostic curve based on mtDNA variant consistency: variant carrier sets are treated as “true” clones, and precision/recall/F1 are computed from the overlap (Jaccard) between variant carrier sets and predicted clades. Panels show example curves for the LARRY and WGS datasets (for comparison to A), as well as for Aged 1 HSC/MPP, MM1, and the MAESTER-CHIP dataset. In all panels, the dashed vertical line indicates the selected cutoff used for downstream analyses (value annotated above each panel).

**Figure S3.**
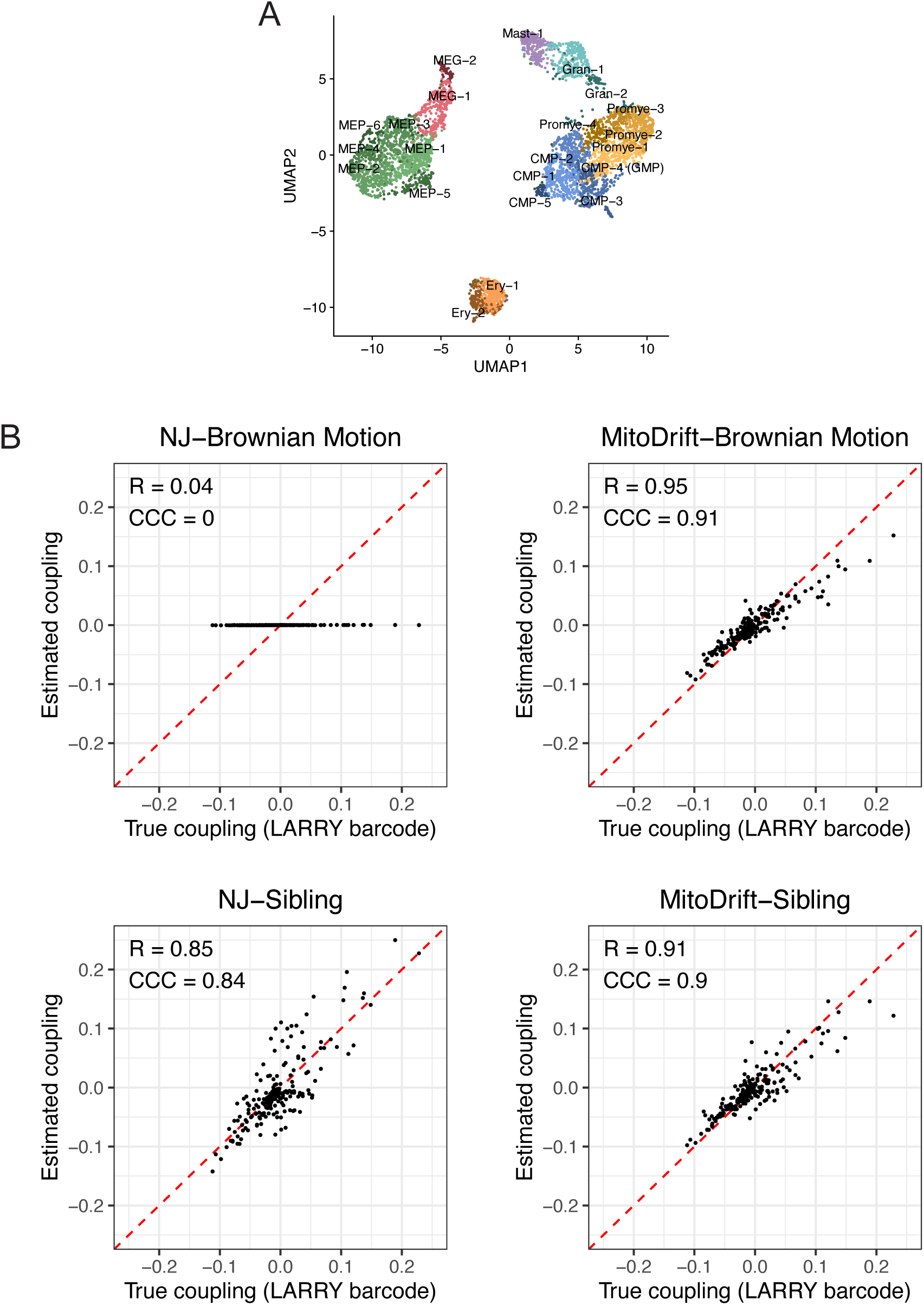
Cell-state map and barcode ground truth validation of coupling estimates. (A) UMAP of the LARRY benchmark with annotated hematopoietic cell states used for lineage-coupling analysis. (B) Agreement between estimated lineage coupling and barcode-defined coupling (“true coupling”) using Neighbor-Joining (NJ) and refined MitoDrift trees, tested in combination with two different phylogenetic distance models: Brownian Motion and Sibling. Dashed lines indicate identity; Pearson’s correlation (R) and Lin’s Concordance Correlation Coefficient (CCC) are shown for each comparison.

**Figure S4.**
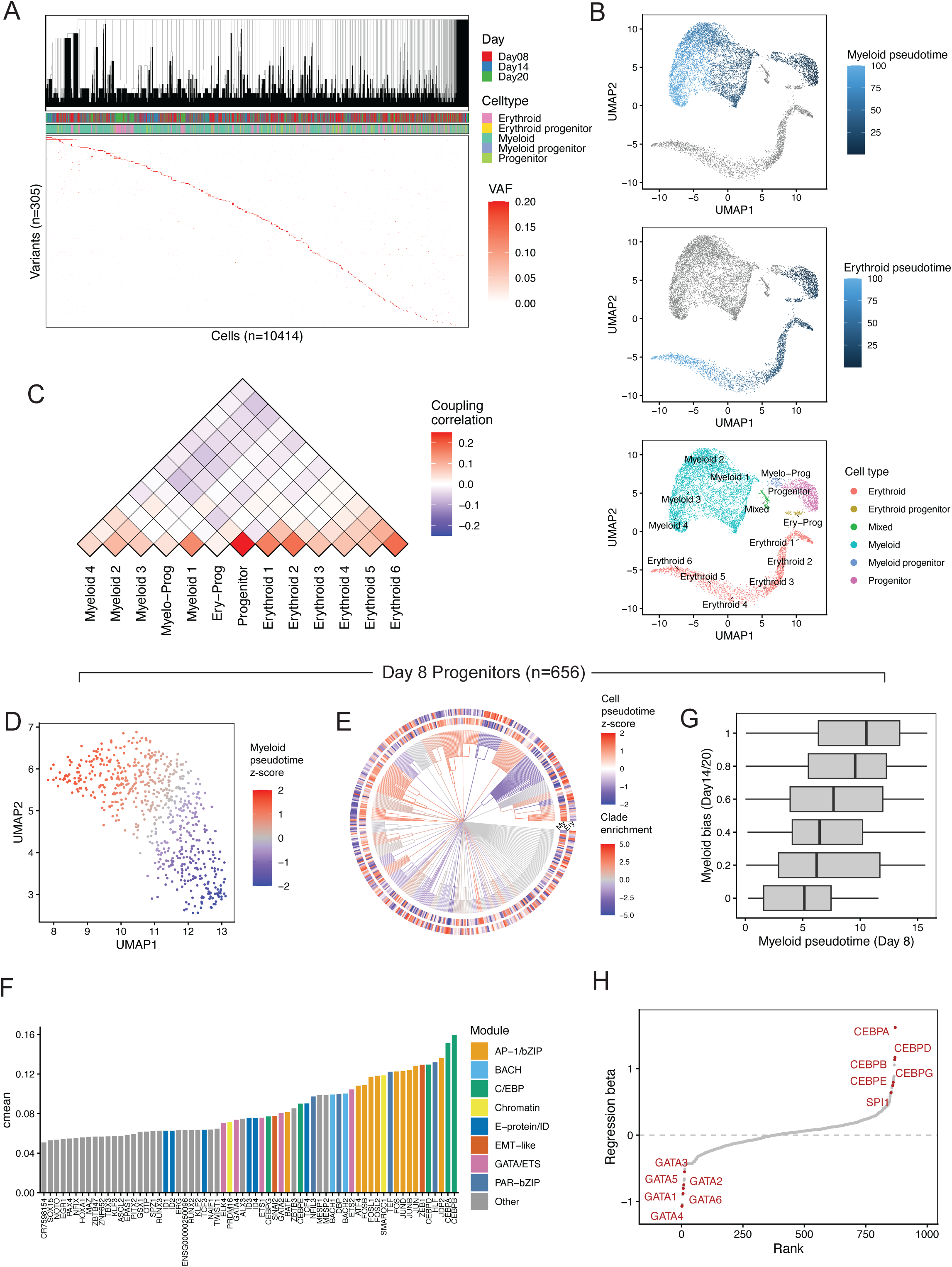
Reanalysis of in vitro CD34+ differentiation mtscATAC-seq data. (A) Cell lineage tree (refined at confidence threshold τ = 0.125) inferred from the Lareau et al. mtscATAC-seq in vitro CD34+ differentiation dataset, juxtaposed with variant allele frequencies (VAF; color scale) across cells (n = 10,414) and mtDNA variants (n = 930); VAF heatmap shows variants detected in ≥15 cells. Cells are ordered by the inferred lineage, with top annotations indicating collection day and annotated cell type. (B) UMAP embedding of the mtscATAC-seq dataset colored by inferred myeloid pseudotime, erythroid pseudotime, and annotated cell types and states. (C) Lineage coupling matrix summarizing pairwise coupling correlations between annotated cell types and states inferred by PATH based on the MitoDrift-inferred lineage tree. (D) UMAP of day 8 progenitors colored by myeloid pseudotime (rescaled z-scores within day 8). (E) MitoDrift lineage tree for day 8 progenitors (refined at confidence threshold τ = 0.125; singletons omitted), with cells annotated by myeloid and erythroid pseudotime (outer ring) and clade-level z-scores for myeloid pseudotime. (F) Phylogenetic signal (cmean) of chromVAR TF and motif activity features within day 8 progenitors, grouped by functional module. (G) Association between day 8 myeloid pseudotime and a myeloid-bias score defined by the fraction of myeloid descendants among day 14 and day 20 cells; boxplots show the distribution of day 8 myeloid pseudotime across bins of the myeloid-bias score. (H) Ranked regression coefficients for TF and motif activity features in day 8 progenitors associated with myeloid-bias at days 14 and 20, with representative regulators highlighted.

**Figure S5.**
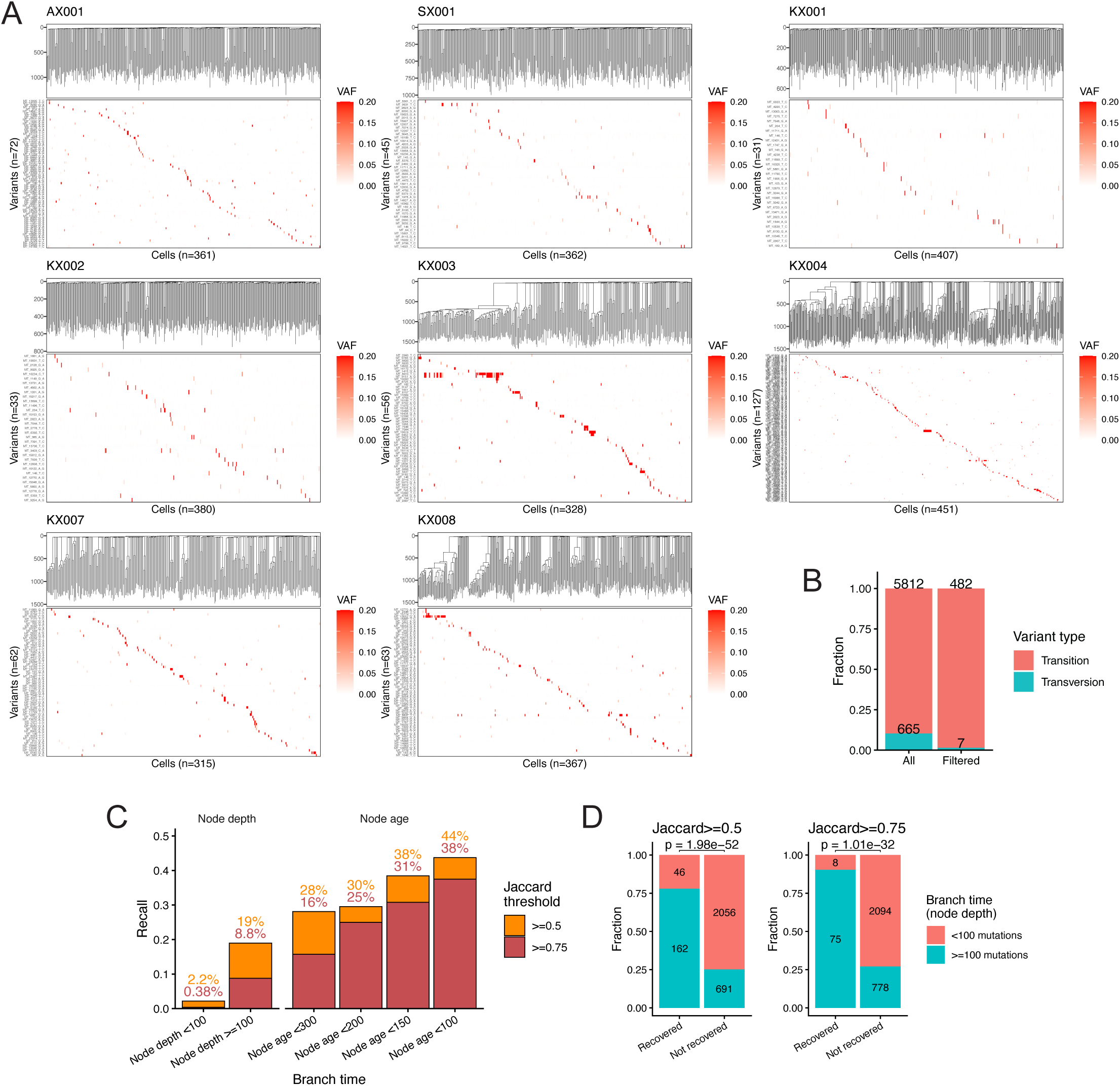
WGS-derived cell lineage trees and mtDNA variants in the in vivo benchmark samples. (A) WGS-derived lineage trees (top) and mtDNA variant allele frequency (VAF) heatmaps (bottom) across single cells for each benchmark sample, with cells ordered by the WGS-derived lineage and mtDNA variants shown on the y-axis (VAF color scale). (B) Fractions of transition and transversion substitutions across all called mtDNA variants and after filtering to variants retained for phylogeny inference. (C) Ground-truth clade recall by MitoDrift (without confidence refinement) as a function of branch time (node depth), evaluated at Jaccard overlap thresholds (≥0.5 and ≥0.75). (D) Fractions of recovered versus not recovered internal nodes at each Jaccard threshold, stratified by branch-time category (<100 versus ≥100 mutations); Fisher’s exact test *P* values are shown above each comparison.

**Figure S6.**
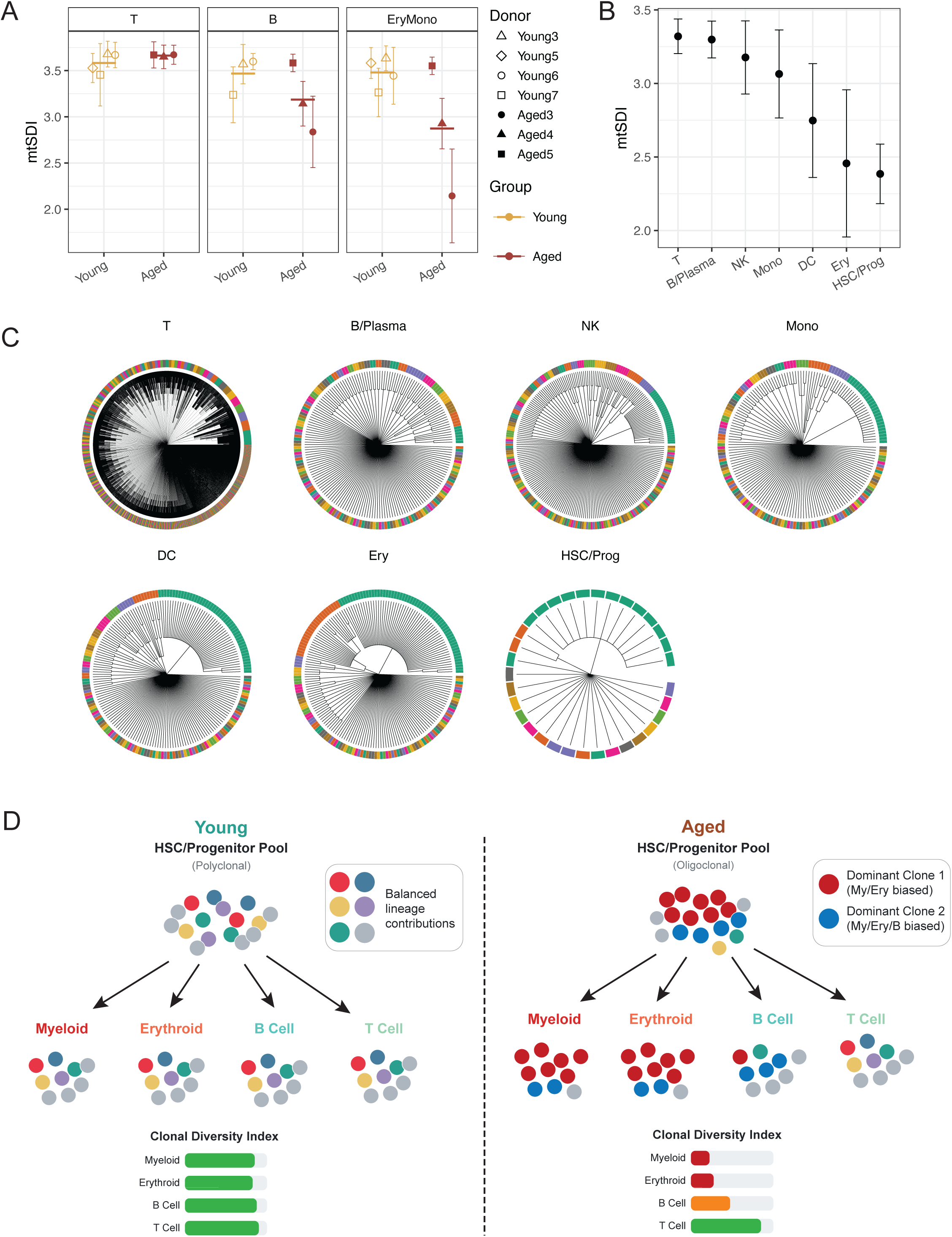
Cell-type-resolved clonal diversity in additional datasets. (A) Shannon diversity index (mtSDI) comparing young versus aged donors across hematopoietic compartments in the donor-multiplexed ReDeeM cohort; computed from 10 random subsamples of 45 cells per cell type per donor (mean ± 95% CI). (B) Mean mtSDI across major hematopoietic compartments in the CHIP donor (Miller et al.) profiled using MAESTER, with error bars representing bootstrap-based 95% confidence intervals. (C) Confidence-refined MitoDrift phylogenies (refined at confidence threshold τ = 0.125) for each hematopoietic cell type in the MAESTER dataset, illustrating cell-type-specific differences in clonal structure. (D) Conceptual schematic contrasting polyclonal versus oligoclonal HSC/progenitor pools and how lineage-biased contributions from dominant clones can differentially reduce clonal diversity across mature lineages.

**Figure S7.**
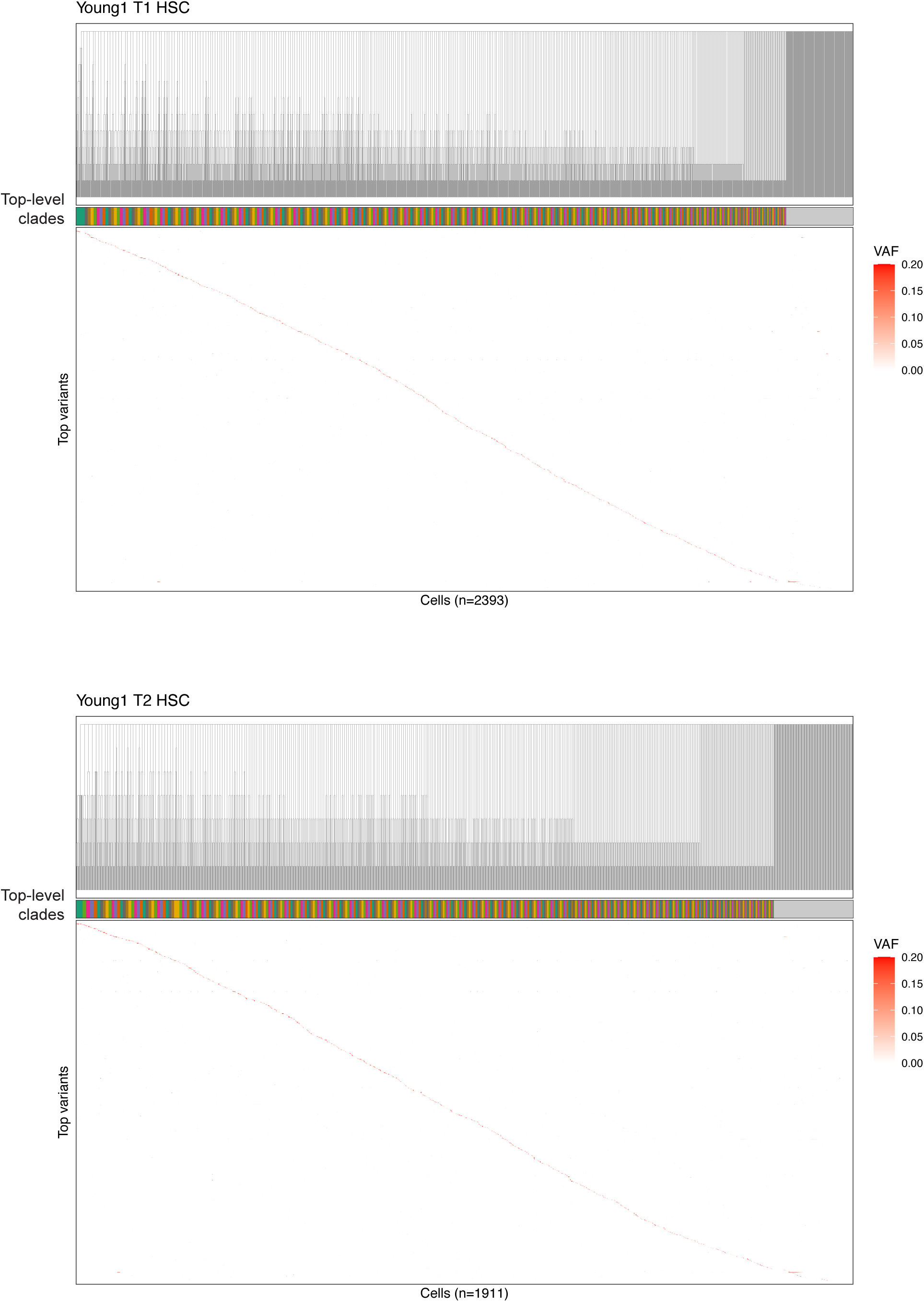
Inferred phylogenies and mtDNA variant patterns in Young1 HSC samples. Heatmaps of mitochondrial variant allele frequency (VAF) across single HSCs from donor Young1 at timepoint T1 (top) and timepoint T2 (bottom). Cells are ordered by the lineage tree inferred by MitoDrift (top dendrogram). The colored annotation bar indicates inferred clade assignments (refined using the expected-misassignment criterion with *ϵ* = 0.002; Methods). Heatmap color denotes per-cell VAF for each mtDNA variant.

**Figure S8.**
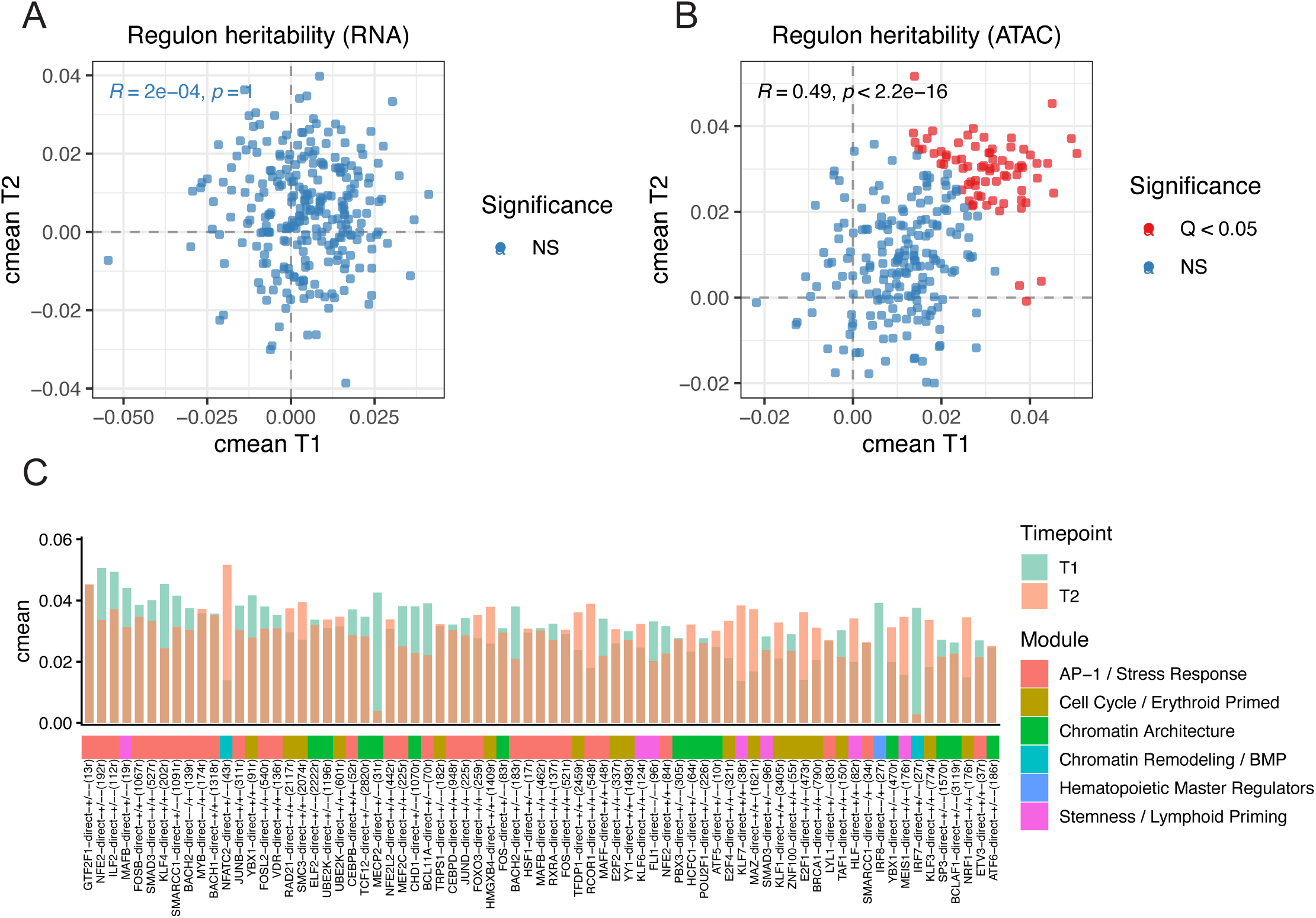
Regulon heritability across timepoints. (A) Scatter plot comparing regulon heritability estimates (cmean) between two longitudinal timepoints (T1 versus T2) using RNA-derived regulon activity scores. (B) Same analysis using ATAC-derived regulon activity scores. Each point denotes a regulon; points are colored by statistical significance in the combined analysis (Q < 0.05). (C) Per-regulon heritability (cmean) at each timepoint for regulons significant in the combined analysis (ATAC-based activity scores), with module assignments indicated.

**Figure S9.**
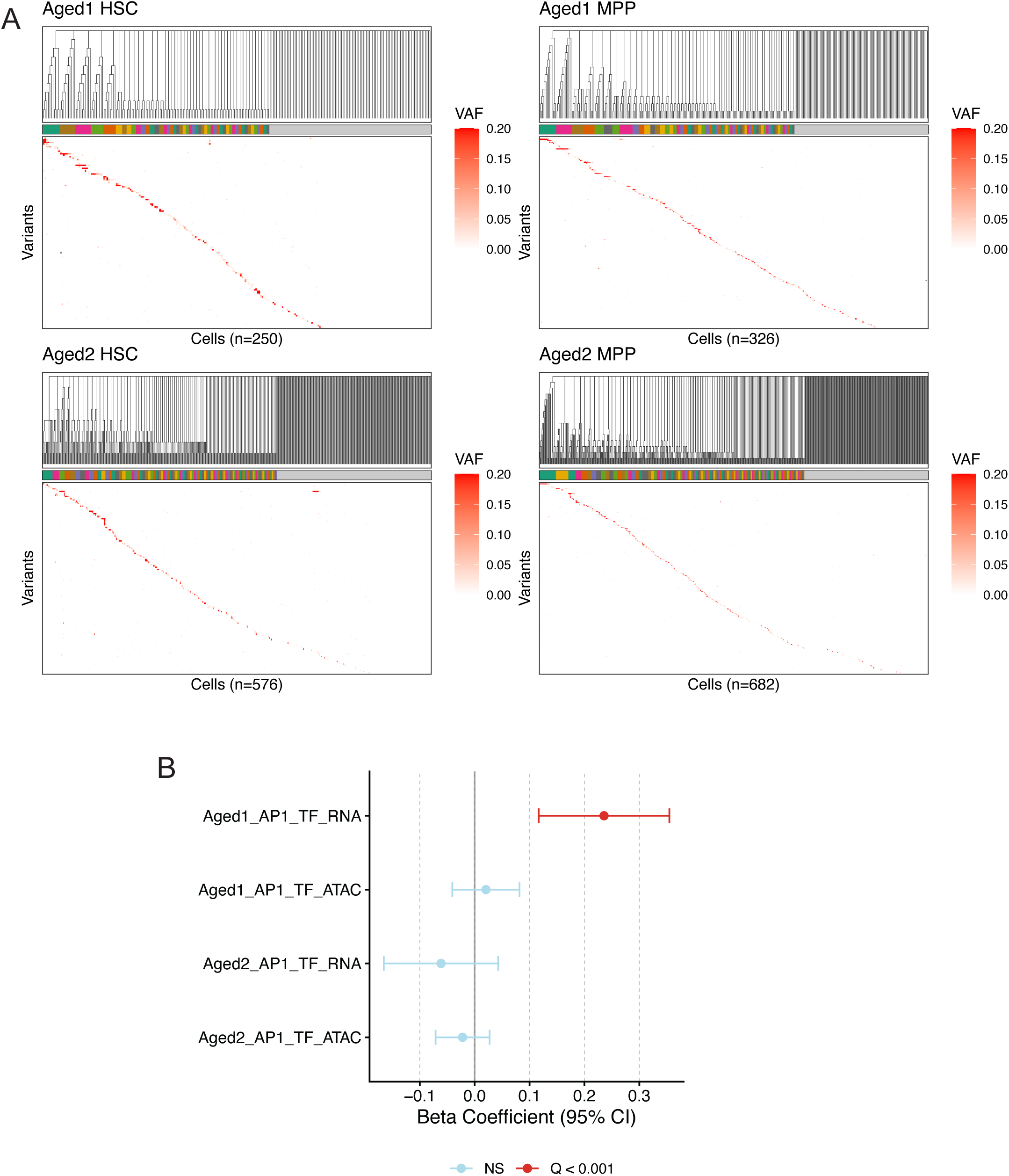
Aged donor HSC/MPP lineage structure and association of AP-1 activity with clonal size. (A) MitoDrift phylogenies (refined at confidence threshold τ = 0.005 for Aged 1, τ = 0.01 for Aged 2) and mtDNA VAF heatmaps for aged donor HSCs and MPPs, with cells ordered by the inferred lineage. Clone assignments (colored bar) are shown above each heatmap. (B) Regression coefficients (beta ± 95% CI) relating average AP-1 TF activity to inferred clone size across aged donors, shown separately for RNA-and ATAC-derived TF activity for each donor.

**Figure S10.**
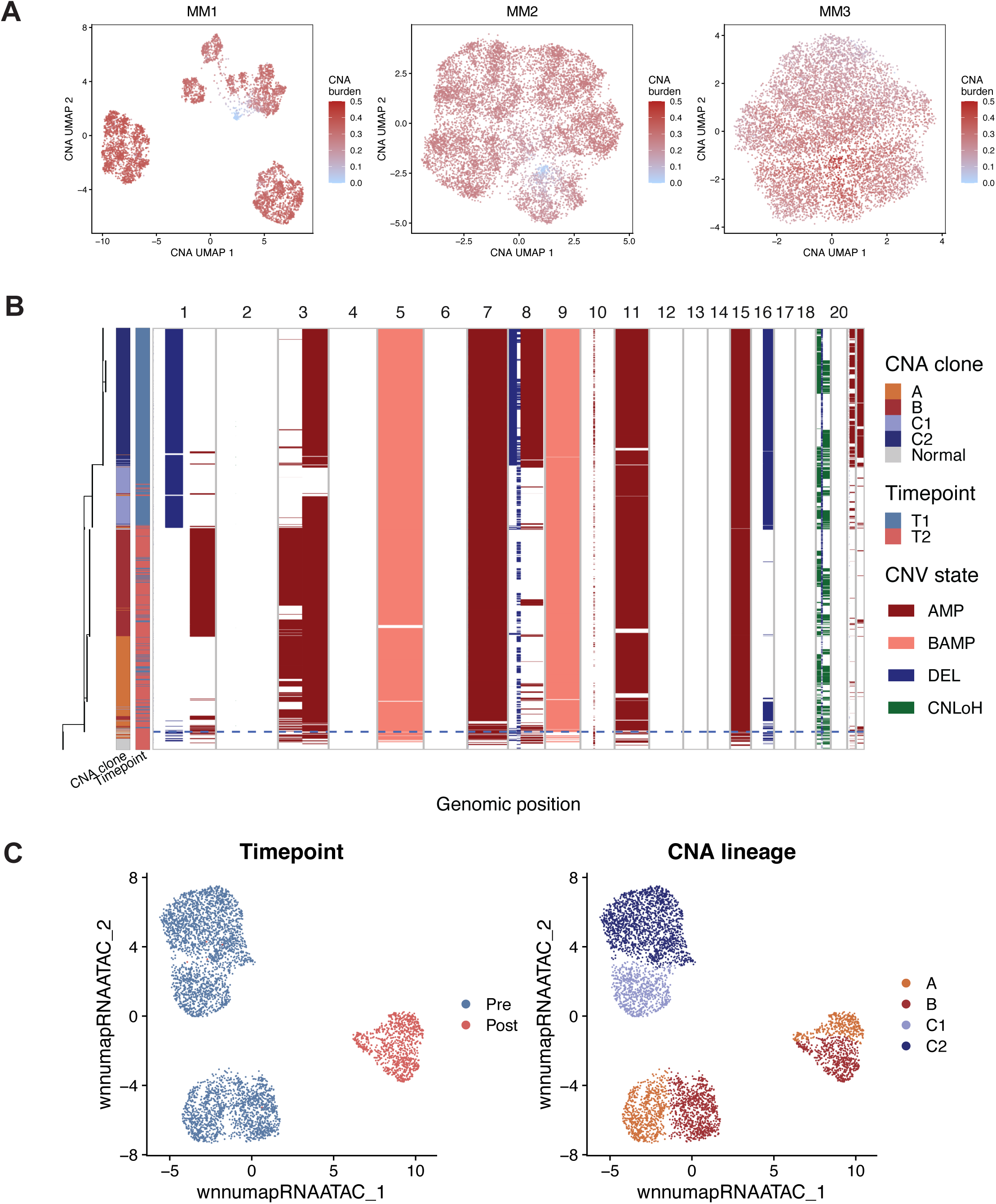
Copy number analysis in myeloma multiome data. (A) CNA-based UMAP projections for plasma cells colored by Numbat-inferred CNA burden for each donor. (B) Heatmap of Numbat-inferred copy-number states for MM1 across the genome for single cells; CNA states are annotated as amplifications (AMP), balanced amplifications (BAMP), deletions (DEL), and copy-neutral loss of heterozygosity (CNLOH). (C) wnnUMAP projection of MM1 malignant plasma cells colored by sampling timepoint and by inferred CNA lineage.

**Figure S11.**
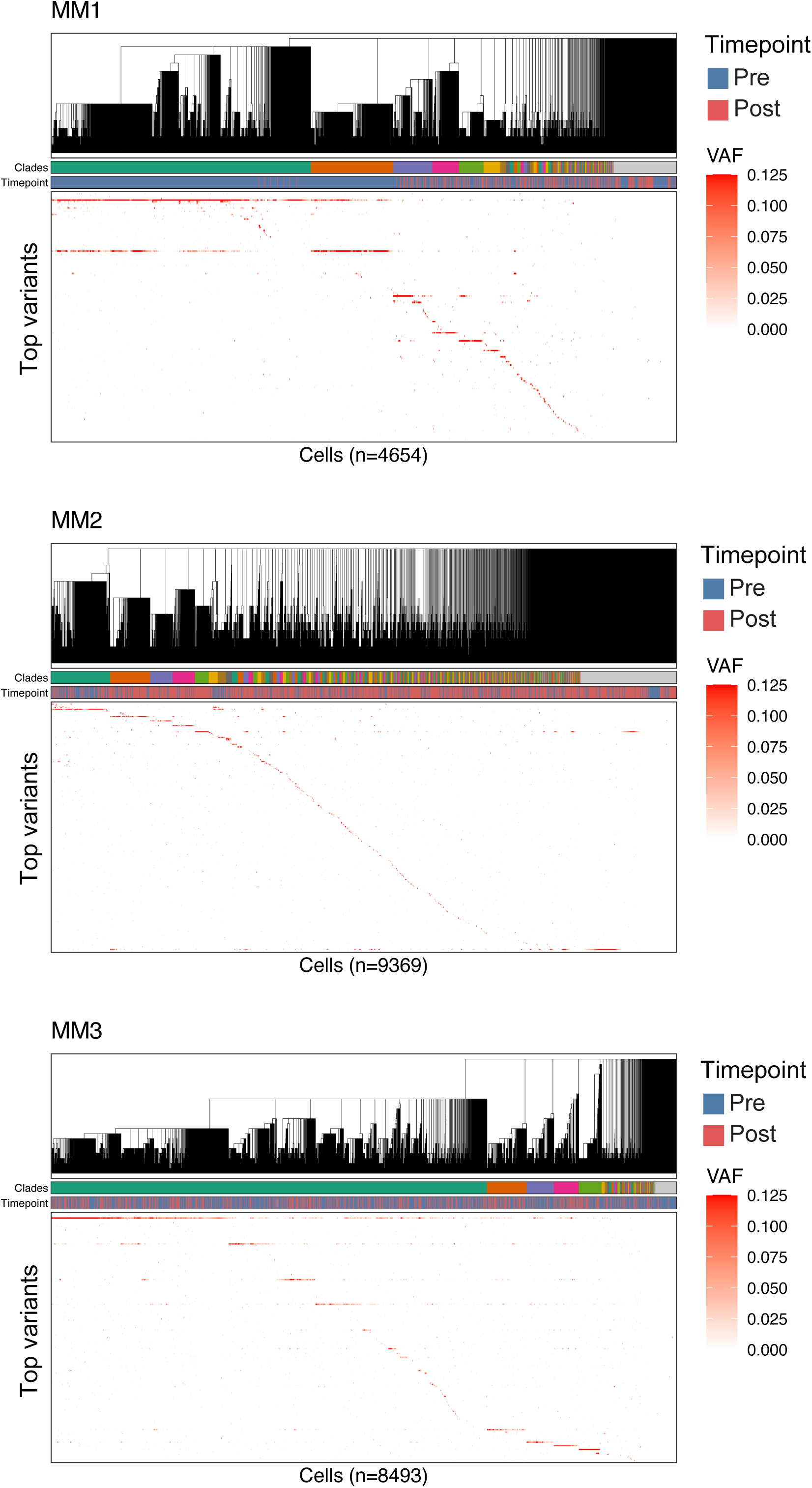
Inferred phylogenies and mtDNA variant patterns in multiple myeloma samples. Heatmaps of mitochondrial variant allele frequency (VAF) across samples from three patients (MM1–3; top to bottom). Cells are ordered by the lineage tree inferred by MitoDrift (top dendrogram; refined at confidence threshold τ = 0.075). The colored annotation bar indicates MitoDrift clone assignments. Heatmap color denotes per-cell VAF for each mtDNA variant.

**Figure S12.**
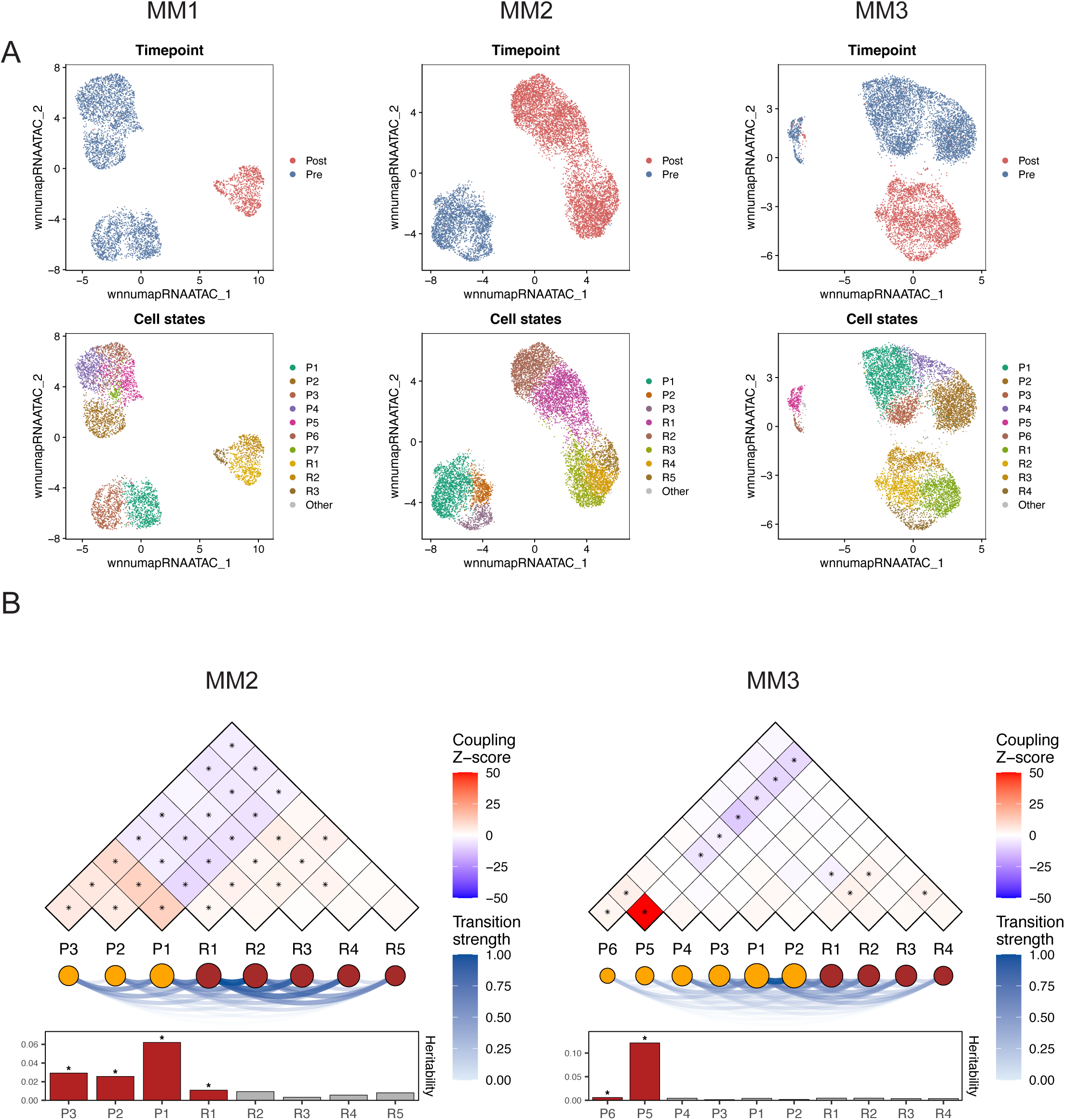
Lineage–state coupling analyses across additional multiple myeloma patients. (A) wnnUMAP projections of malignant plasma cells from three multiple myeloma patients (MM1–MM3) colored by sampling timepoint (pre- versus post-treatment; top) and by annotated malignant cell states (bottom). (B) Cell state coupling summary for MM2 and MM3, showing lineage coupling between malignant cell states (pairwise coupling Z-scores), inferred transition strengths between states, and per-state heritability on the cell lineage tree.

**Figure S13.**
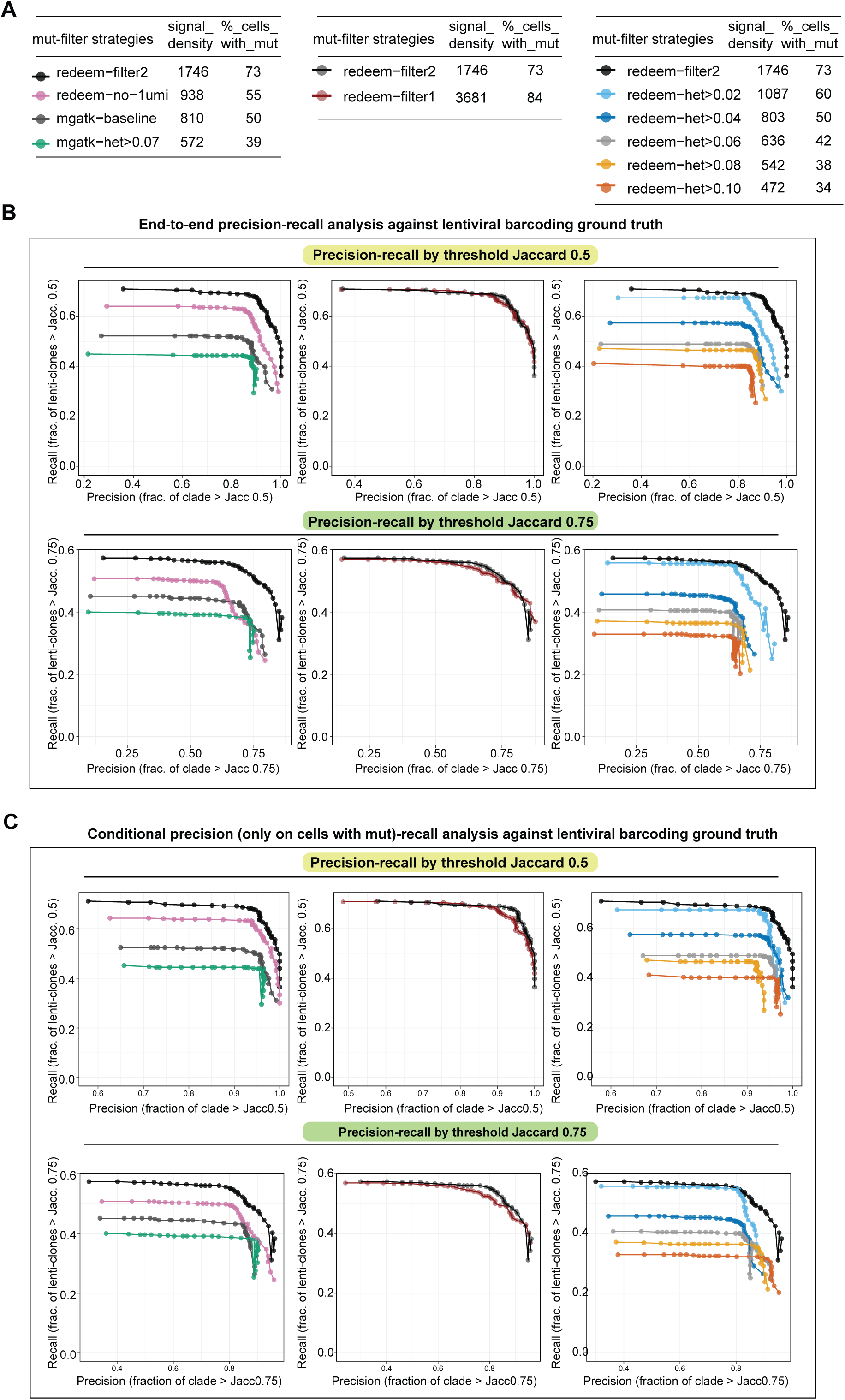
Benchmarking the impact of mtDNA variant filtering on lineage reconstruction performance. (A) Summary of mutation retention after applying each predefined variant-filtering strategy. Signal_density reports the number of retained mutation observations per 1,000 cells after filtering, where a mutation observation is a retained cell–variant entry. %_cells_with_mut reports the percentage of cells retaining ≥1 mutation after filtering (i.e., evaluable cells). (B) End-to-end precision–recall benchmarking of phylogenetic reconstruction across filtering strategies, evaluated against lentiviral barcoding (LARRY) ground-truth clone labels (clone size ≥2) at Jaccard overlap thresholds of 0.5 and 0.75. The three comparison groups (left to right) are (i) ReDeeM vs mgatk preprocessing, where ReDeeM is evaluated under ReDeeM filter-2 and ReDeeM filter-2 with 1UMI-supported mutations removed, and mgatk is evaluated under its baseline settings and an additional heteroplasmy ≥0.07 filtering regime recommended in prior work (Lareau et al., 2021, Nat. Biotech.); (ii) ReDeeM filter-1 vs filter-2; and (iii) ReDeeM filter-2 under increasing heteroplasmy thresholds. Precision and recall are computed on the same full set of cells for each condition (“end-to-end”), thus capturing both clade accuracy and signal dropout effects. (C) Conditional-precision analysis of precision–recall using the same benchmarking framework as in (B), but computing precision only among evaluable cells (cells retaining ≥1 mutation observation after filtering) while retaining full-cell recall. This isolates clade accuracy among retained-signal cells from mutation dropout effects. Across panels (B–C), inferred clades are obtained by collapsing the MitoDrift phylogeny across a sweep of confidence refinement thresholds (τ). Precision is defined as the fraction of inferred clades whose best-matching ground-truth clone exceeds the indicated Jaccard threshold, and recall is defined as the fraction of ground-truth clones whose best-matching inferred clade exceeds the threshold. Curves are averaged across 10 matched clone-based subsets reused across all conditions to control for subset composition effects, with matched inference parameters across conditions.

**Figure S14.**
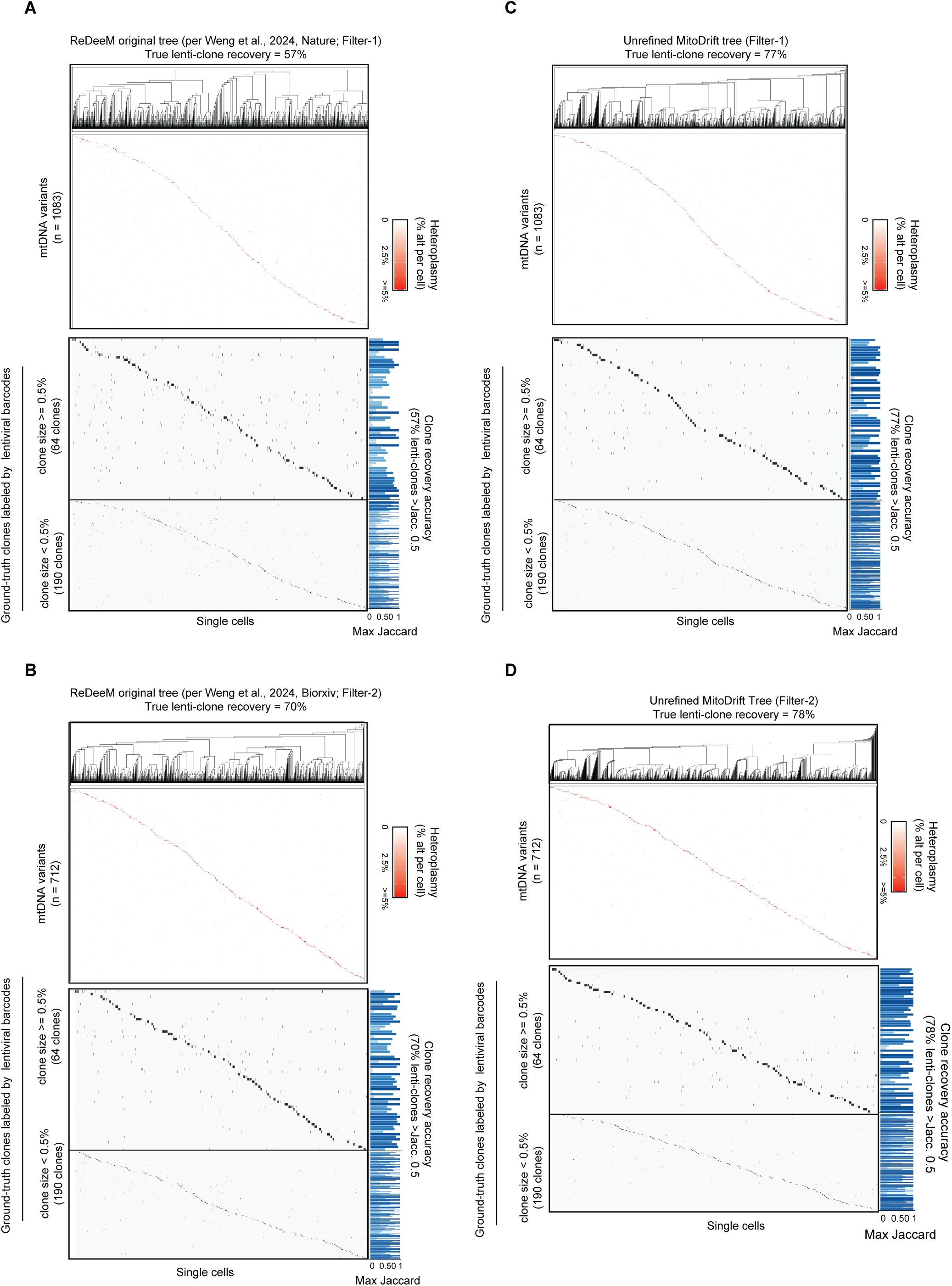
Ground-truth clone recovery in original ReDeeM and unrefined MitoDrift trees. (A–B) Original ReDeeM tree (Neighbor-Joining on binarized variants). Phylogeny constructed using the ReDeeM workflow: mtDNA variants binarized (presence/absence), pairwise weighted Jaccard distances computed, and Neighbor-Joining (NJ) tree inferred. (A) Filter-1 (Weng et al., 2024, Nature). (B) Filter-2 (stricter filtering; Weng et al., 2024, bioRxiv). (C–D) Unrefined MitoDrift tree (NJ backbone, pre-calibration). Phylogeny represents the NJ backbone used in MitoDrift before posterior-based clade collapsing. The workflow is: mtDNA variants are not binarized; Manhattan distances are computed from continuous heteroplasmy profiles, followed by NJ inference. (C) Filter-1. (D) Filter-2. Each panel displays: top, inferred NJ phylogeny; middle, mtDNA variant heteroplasmy heatmap (rows, variants; columns, cells ordered by tree; color, alternate-allele fraction); bottom, ground-truth lentiviral clones stratified by size (≥0.5%, 64 clones; <0.5%, 190 clones). Clone recovery accuracy is defined as the fraction of lentiviral clones whose best-matching clade achieves Jaccard overlap >0.5. Filter-2 improves recovery versus Filter-1. The heteroplasmy-based unrefined MitoDrift tree (Manhattan distance) outperforms the binarized ReDeeM tree (weighted Jaccard) given the same mutation filtering strategy.

**Table S1.** MitoDrift run parameters used across analyses. Summary of MitoDrift run parameters (datasets, samples included, MCMC iterations, ASDSF threshold, number of chains, and burn-in) used across analyses and figure panels.

| Analysis | Dataset | Samples included | MCMC iter | ASDSF thres | Chains | Burn-in |
| --- | --- | --- | --- | --- | --- | --- |
| pLARRY tree accuracy benchmark (full tree) | pLARRY | pL1000 | 150000 |  | 50 | 10000 |
| pLARRY tree accuracy benchmark (subsample trees) | pLARRY | pL1000 (subset seeds 1–10) |  | 0.05 | 50 | 2000 |
| WGS benchmark (subsampled trees) | WGS benchmark (Mitchell et al.) | AX001, SX001, KX001–KX004, KX007–KX008 (seeds 1–10) | 10000 |  | 50 | 1000 |
| WGS benchmark (full donor trees) | WGS benchmark (Mitchell et al.) | AX001, SX001, KX001–KX004, KX007–KX008 | 20000 |  | 20 | 2000 |
| MAESTER-CHIP benchmark | MAESTER-CHIP (Miller et al.) | MAESTER-CHIP |  | 0.1 | 10 | 10000 |
| Extended ReDeeM cohort mtSDI by lineage (cell-type subsamples) | ReDeeM | Young1, Young2, Young8, Aged1, Aged2 x cell types (seeds 1–20) | 35000 |  | 20 | 5000 |
| Multiplexed cohort mtSDI by lineage (hash-refined subsamples) | ReDeeM (multiplexed) | BMMC2, BMMC3, BMMC8, BMMC10, SBM1253, SBM1028, SBM1037 x cell types (seeds 1–10) | 5000 |  | 20 | 500 |
| AP-1 in aged HSC and MPP | ReDeeM | Aged1, Aged2 (HSC, MPP) |  | 0.05 | 50 | 4000 |
| Young1 longitudinal HSC regulon heritability (refined HSC trees) | ReDeeM | Young1 T1, T2 (refined HSC) |  | 0.1 | 20 | 10000 |
| Myeloma joint-timepoint phylogenies | Myeloma multiome | MM1 plasma (all timepoints) |  | 0.1 | 20 | 10000 |
|  | Myeloma multiome | MM2 plasma (all timepoints) |  | 0.1 | 10 | 10000 |
|  | Myeloma multiome | MM3 plasma (all timepoints) |  | 0.1 | 10 | 10000 |
| In vitro CD34+ differentiation (800-cell subset) | CD34+ differentiation (Lareau et al.) | mtscATAC | 350000 |  | 10 | 35000 |

